# Broad kinase inhibition mitigates early neuronal dysfunction and cognitive deficits in tauopathy

**DOI:** 10.1101/2020.07.31.229583

**Authors:** Shon A. Koren, Matthew J. Hamm, Ryan Cloyd, Sarah N. Fontaine, Emad Chishti, Chiara Lanzillotta, Jennifer Rodriguez-Rivera, Alexandria Ingram, Michelle Bell, Sara M. Galvis-Escobar, Nicholas Zulia, Fabio Di Domenico, Duc Duong, Nicholas T. Seyfried, David Powell, Moriel Vandsburger, Tal Frolinger, Anika M.S. Hartz, John Koren, Jeffrey M. Axten, Nicholas J. Laping, Jose F. Abisambra

## Abstract

Tauopathies are a group of more than twenty known disorders that involve progressive neurodegeneration, cognitive decline, and pathological tau accumulation. Current therapeutic strategies provide only limited, late-stage symptomatic treatment. This is partly due to lack of understanding of the molecular mechanisms linking tau and cellular dysfunction, especially during the early stages of disease progression. In this study, we treated early stage tau transgenic mice with a multi-target kinase inhibitor to identify novel substrates that contribute to cognitive impairment and exhibit therapeutic potential. Drug treatment significantly ameliorated brain atrophy and cognitive function as determined by behavioral testing and a sensitive imaging technique called manganese-enhanced magnetic resonance imaging (MEMRI) with quantitative R1 mapping. Surprisingly, these benefits occurred despite unchanged hyperphosphorylated tau levels. To elucidate the mechanism behind these improved cognitive outcomes, we performed quantitative proteomics to determine the altered protein network during this early stage in tauopathy and compare this model with the human AD proteome. We identified a cluster of preserved pathways shared with human tauopathy with striking potential for broad multi-target kinase intervention. We further report high confidence candidate proteins as novel therapeutically relevant targets for the treatment of tauopathy.

**One Sentence Summary:** Multi-target kinase inhibition rescues cognitive function in early stage tauopathy mice and reverses proteomic shifts common to Alzheimer’s disease in humans.

## Introduction

Tauopathies, the most common of which is Alzheimer’s disease (AD), are a group of neurological disorders defined by the neuropathological accumulation of tau protein that present progressive cognitive dysfunction and brain atrophy. No cure for tauopathies exists, and current treatment strategies are palliative (*1, 2*). Several clinical trials have targeted toxic tau species through immunotherapies or by inhibiting tau post-translational modifications and fibrillization with mixed results (*1–4*). However, the disease etiology of tauopathies, AD in particular, are almost certainly multifactorial. A promising strategy to mitigate the complex nature of tauopathies, similar to established approaches for other chronic, progressive diseases, encompass multi-target or combinatorial treatments (*3, 5*). These strategies are limited by requiring a thorough understanding of the cellular perturbations that underly the disease.

Modern systems biology approaches provide unparalleled power in investigating the cellular alterations in disease. These tools highlight promising candidate targets for novel therapeutics while elucidating the pathophysiology of the disease. Recent studies focused on the molecular network changes in AD suggest concurrent changes in energy metabolism, immune response, synapse activity, cytoskeletal stability, and RNA metabolism pathways present early in disease progression (*6–8*). These reports validate decades of research utilizing *in vitro* and *in vivo* disease models. Using these tools, the pathways most closely associated with early cognitive decline and neurodegeneration can be determined and used to assess novel therapeutics. Importantly, these resources have not yet been applied to establish the efficacy of multi-target drugs in mitigating cognitive decline in animal models of tauopathy.

Aberrant phosphorylation and kinase signaling is a hallmark of AD and other tauopathies (*8, 9*). Here, we report the use of a multi-target kinase inhibitor, GSK2606414 or 414, to treat tau transgenic mice in a proof-of-concept study to evaluate whether such strategies can mitigate the early negative outcomes of tauopathy. This compound potently inhibits multiple tyrosine and serine/threonine kinases at low micromolar concentrations (*10*) and ameliorates phenotypes in a variety of neurological disorder and neurodegenerative models (*11–16*). The targets of 414 include kinases involved in tauopathy pathogenesis, including protein kinase R-like endoplasmic reticulum kinase, or PERK (*10–12*), the MAPK cascade (*10*), receptor-interacting serine/threonine-protein kinase 1, or RIPK1 (*17*), and KIT (*18*). We show that brain atrophy and phenotypes of cognitive dysfunction evident in this mouse model of tauopathy are substantially reduced by treatment with 414. Using manganese-enhanced MRI (MEMRI) with parametric mapping, we show that multi-target kinase inhibition rescues deficits in hippocampal calcium activity. The rescue of tauopathy occurs in the absence of changes to toxic, hyper-phosphorylated tau levels and without inhibition of PERK, the target canonically associated with the neuroprotective benefits of 414 (*10, 11*). We use quantitative proteomics to investigate both the cellular alterations and the networks responsive to multi-target kinase inhibition associated with cognitive rescue in this early stage tauopathic mouse model. Finally, we identify novel candidate proteins that are rescued by drug treatment and are consistent between the tauopathy mouse and human proteome. These analyses identify novel candidate targets for therapeutic intervention in future studies.

## Results

### Tau transgenic mice exhibit common neurodegenerative features ameliorated by multi-target kinase inhibition

To investigate the pathways responsible for early cognitive impairment in tauopathy and their amenability to treatment with a multi-target kinase inhibitor, we used the rTg4510 tau transgenic mouse model of frontotemporal lobar degeneration (FTLD). The rTg4510 (Tg) mice exhibit well-characterized disease progression driven by the overexpression of P301L human tau primarily in forebrain neurons (*19–21*). Neurofibrillary tau pathology appears as early as two months, with progressively worsening cognitive impairment detectable as early as two and a half months. Additionally, significant brain atrophy is detectable in this model by 5 months of age (Fig. 1A).

**Fig 1:**
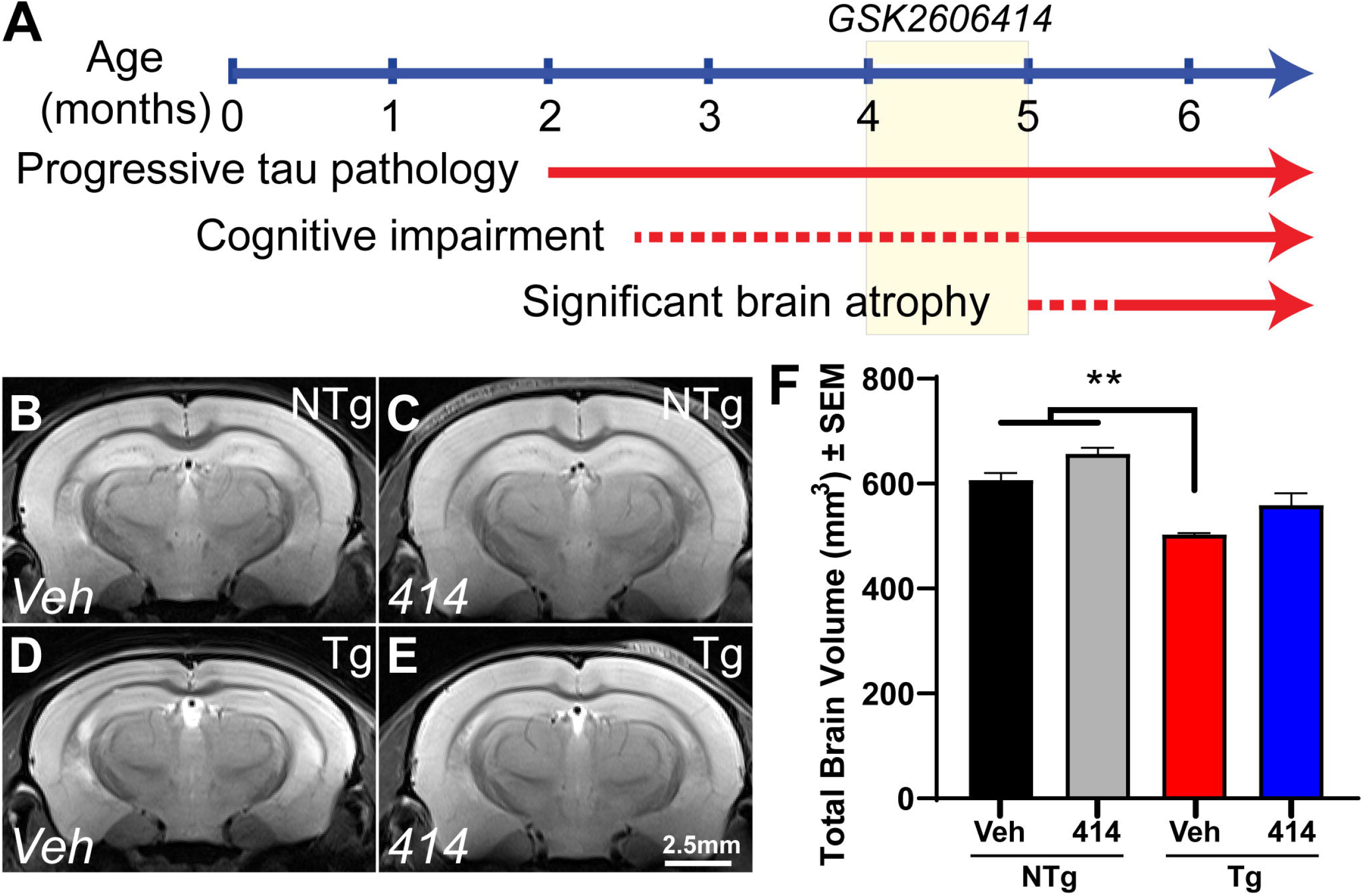
Multi-target kinase inhibition with GSK2606414 rescues brain atrophy in rTg4510 tau transgenic mice. **A** Timeline of tauopathy phenotype in transgenic (Tg) mice and depiction of drug delivery between 4-5mo. **B-E** Representative T2-weighted MR images of non-transgenic (NTg) and Tg mice treated with vehicle (Veh) or GSK2606414 (414). **F** Total brain volume quantification (NTg + 414, n = 2; others n = 3). Two-way ANOVA with Tukey *post hoc* test. Data are expressed as the mean ± SEM, **p < 0.01.

We treated four-month-old Tg and non-transgenic control (NTg) mice with the multi-target kinase inhibitor GSK2606414 (414) for 30 days. Four-month-old Tg mice display significant tau burden, but they do not present with severe cognitive impairment and therefore provide a therapeutic window amenable to treatment strategies. T2-weighted MRI (Fig. 1B-F) revealed that Tg mice treated with vehicle (Tg + Veh) exhibited an approximately 20% reduction in total brain volume compared with non-transgenic control mice treated with either vehicle (NTg + Veh, **p = 0.0063) or 414 (NTg + 414, **p = 0.0012). Surprisingly, tau transgenic mice treated with 414 (Tg + 414) exhibited only a 10% reduction in total brain volume that lacked statistical significance from NTg + Veh control animals (ns, p > 0.15).

We previously established that this tau transgenic mouse line exhibits altered calcium homeostasis detected by MEMRI with R1 mapping as early as 3mo of age which continues as the mice age (*22*). Here, we corroborated those findings, identifying altered ΔR1 in 5mo Tg mice (Fig. 2A-D). The strongest differences in ΔR1 observed were found in hippocampal regions such as the dentate gyrus (DG, Fig. 2E) and Cornu ammonis area 1 (CA1, Fig. 2F). In both regions, our experimental Tg + Veh mice exhibited significantly decreased ΔR1 when compared to the control NTg + Veh mice (DG, ***p = 0.0003; CA1, *p = 0.0226). Tg + 414 treated mice displayed no apparent impairment in calcium activity measured by MEMRI in both regions relative to NTg + Veh mice (DG and CA1, ns) and displayed significant improvement in calcium activity relative to the Tg + Veh animals (DG, **p = 0.0019; CA1, *p = 0.0211). These findings demonstrate that, in tandem with the changes in total brain volume, treatment with the multi-target kinase inhibitor 414 rescued hippocampal activity deficits in this model.

**Fig 2:**
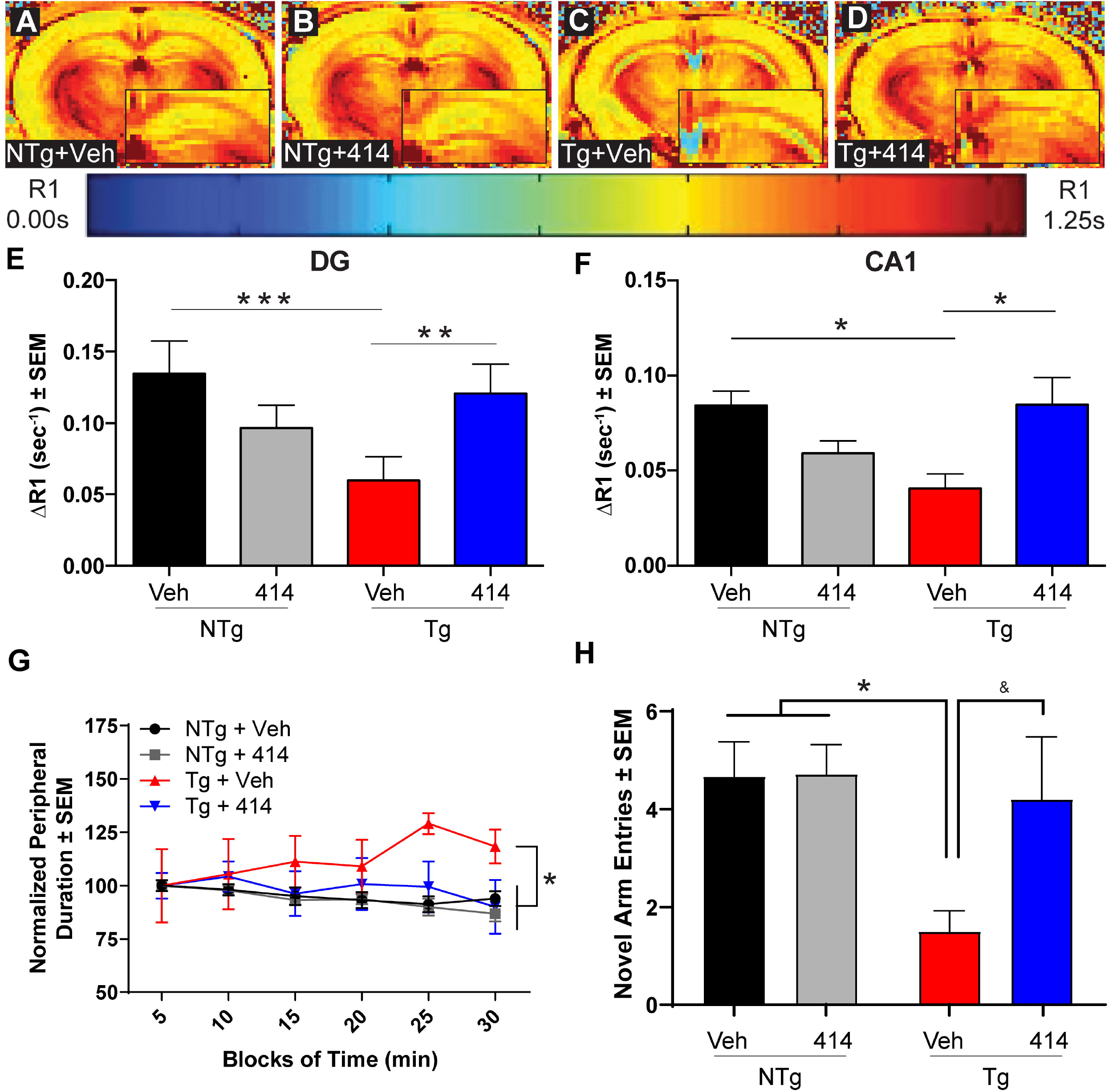
GSK2606414 treatment rescues brain dysfunction in tau transgenic mice. **A-D** Representative DR1-rendered MEMRI images. The insets in panels A-D show half of the hippocampus of each representative image. **E-F** Calculated DR1 values in the dentate gyrus (DG, **E**) and Cannu ammonis 1 (CA1, **F**). Two-way ANOVA with Tukey *post hoc* test. Data are expressed as the mean ± SEM, *\*p* < 0.05, *\*\*p* < 0.01, *\*\*\*p* < 0.001, (n = 4-5). **G** Results of open field behavioral task indicating percent of time spent in the periphery, normalized to the first five minutes (n = 6-10). Two-way ANOVA with Tukey *post hoc* test. Data are expressed as the mean relative to NTg + Veh ± SEM, *\*p* < 0.05. **H** Y-maze behavioral task results for number of novel arm entries (n = 5-7). Two-way ANOVA with Tukey *post hoc* test. Data expressed as the mean ± SEM, *\*p* < 0.05, & denotes p = 0.1095.

Since calcium dyshomeostasis underlies cognitive impairments in tauopathy (*23*), we evaluated the effects of 414 in two behavioral paradigms: open field as a measure of anxiety-like phenotype, and Y-maze as a measure of cognition. The open field behavioral paradigm demonstrated a Tg phenotype wherein Tg + Veh mice spent more time in the field periphery than either NTg group (NTg + Veh: **p = 0.0012; NTg + 414: ***p = 0.0001), suggesting a transgene-dependent anxiety-like behavior (Fig. 2G). Tg mice treated with 414 spent significantly less time in the periphery than Veh treated Tg mice (*p=0.0260) and were statistically indistinguishable from either NTg treatment group (ns, p > 0.15). In the Y-maze, Tg + Veh mice exhibited decreased novel arm entries (*p = 0.0365) compared to NTg control mice. Treatment of Tg animals with 414, though statistically indistinguishable from the NTg groups (ns, p > 0.15), only partially rescued the number of entries (ns, p = 0.1095) compared to vehicle-treated Tg mice (Fig. 2H). Total time spent in the novel arm indicated Tg + Veh mice had reduced performance compared to NTg + Veh mice (*p = 0.0433). Again, the Tg + 414 treatment group was statistically indistinguishable from either NTg group (ns, p > 0.15) but displayed an incomplete rescue of this phenotype as compared to Tg + Veh mice (ns, p = 0.0882) (Supplemental Fig. 1). Consistent with our MEMRI and brain volume data, these data demonstrated that treatment with 414 mitigates the cognitive deficits of early-stage tauopathy in this model.

### GSK2606414 rescues functional deficits without altering tau hyper-phosphorylation

A previous study reported that GSK2606414 reduced tau pathology by preventing PERK activation and consequent enhancement of GSK3β activity (*11*). GSK3β phosphorylates tau at sites that are associated with tauopathy progression (*24*). To test whether decreased tau phosphorylation was responsible for the cognitive rescue in Tg mice, we measured the levels of disease-associated, hyper-phosphorylated tau at the S396/S404 (PHF1) and total tau levels (Fig. 3A). 414 did not modify the levels of PHF1, total tau, or the relative phosphorylation at the PHF1 epitope (Fig. 3B-D).

**Fig 3.**
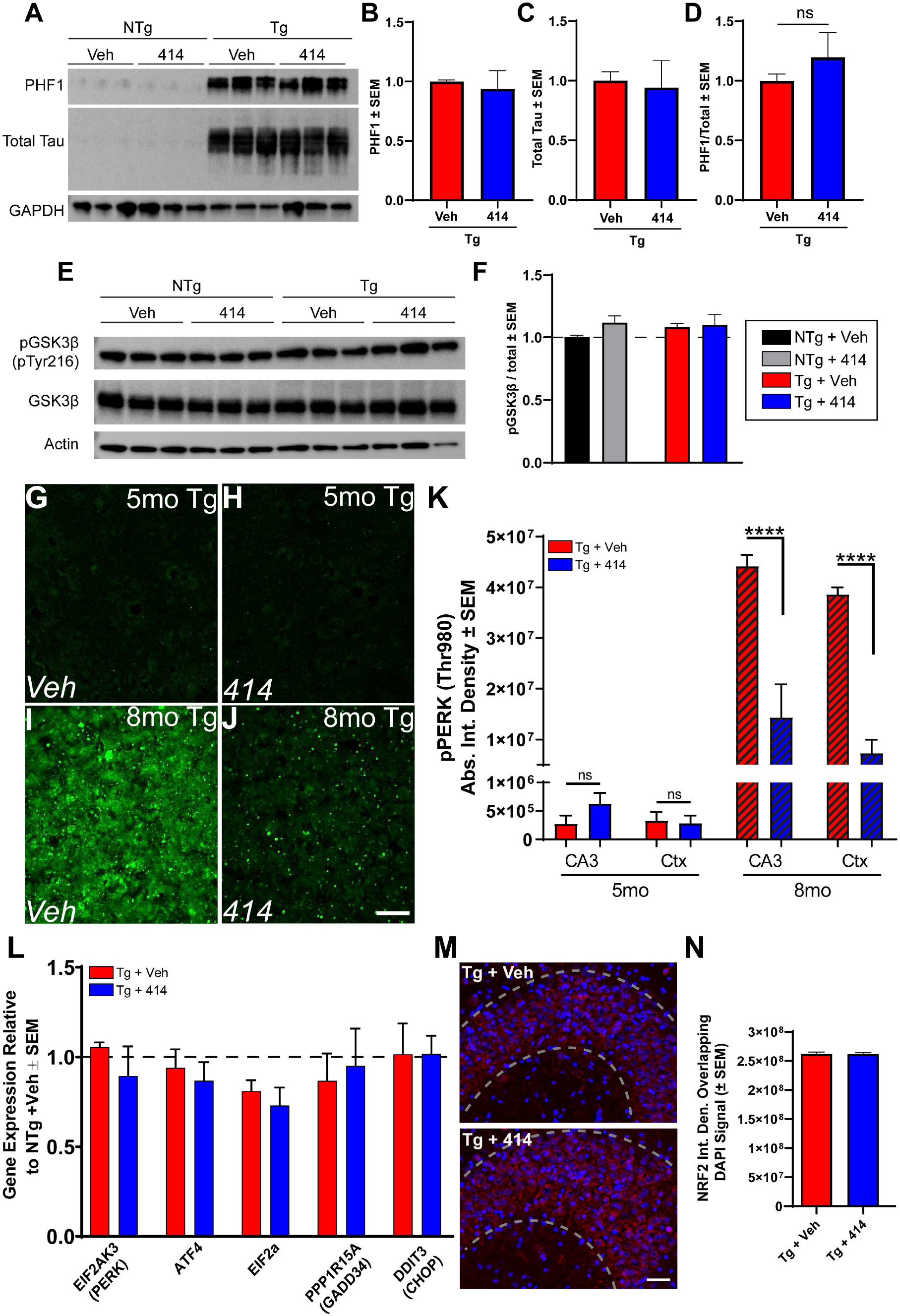
GSK2606414-mediated cognitive rescue in early-stage tau transgenic mice is independent of tau hyper-phosphorylation and GSK3β activity. **A** Immunoblots of PHF1 (S396/S414) and total tau in 5mo NTg and Tg animals treated with vehicle or GSK2606414 (414). **B-D** Results of PHF1 (**B**), total tau (**C**), and relative phosphorylation (**D**) is normalized to GAPDH (n = 3). Two-way ANOVA with Tukey *post hoc* test. Data are expressed as the mean relative to Tg + Veh ± SEM. **G-J** Representative images of active, pT980 PERK (pPERK) staining. All images were post-processed equally and highlights lack of positive staining in 5mo Tg mice. Scale bar = 25 microns. **K** Quantification of pPERK staining reveals that 414 treatment has no detectable change in 5mo Tg mice (n = 5-7), but strongly inhibits PERK activity in the CA3 and superior medial cortex (Ctx) in 8mo Tg mice (hatched bars, n = 3). Two-way ANOVA with Sidak *post hoc* test. Data are expressed as the mean relative to NTg + Veh ± SEM, *\*\*\*\*p* < 0.0001. **L** Relative hippocampal gene expression of direct mediators of the PERK-UPR pathway in 5mo NTg and Tg mice treated for 30d with Veh or 414 normalized to GAPDH and 18S (n = 4-6). Two-way ANOVA with Tukey *post hoc* test. Data are expressed as the mean relative to NTg + Veh ± SEM, *p < 0.05, **p < 0.01. **M-N** Representative images (**M**) and quantification (**N**) of overlap between NRF2 (red) and nuclear staining (DAPI, blue) in the CA3 of 5mo NTg and Tg mice treated with either Veh or 414 (n = 3). Unpaired Student’s t-test. Error bars denote SEM. Scale bar = 25 microns.

### Functional deficits of early stage tau transgenic mice do not depend on PERK/UPR activation

GSK2606414 was developed as a first-in-class inhibitor of PERK (*10*). PERK, an ER-associated kinase, functions as part of the unfolded protein response (UPR) by attenuating global translation and dampening the level of incoming nascent proteins in the ER under conditions of ER stress (*25*). However, long term PERK activation leads to sustained suppression of translation and activation of pro-apoptotic signaling cascades. Previous studies have used 414 to inhibit PERK to prevent neurodegeneration by limiting the deleterious chronic activation of the UPR (*11, 12, 25*). We evaluated whether this PERK activity was responsible for neurodegeneration in early stage Tg mice. Consistent with previous reports (*11, 26, 27*), PERK activation (as measured by the phosphorylation of T980) was minimally detected by immunofluorescent stains of the hippocampus and cortex of 5mo Tg mice (Fig. 3G-H). At 8mo, Tg mice had over one hundred-fold greater levels of phosphorylated PERK (Fig. 3I-J) which was inhibited by 414 treatment as previously reported (Fig. 3K) (*11*). Quantitative real-time PCR gene card arrays verified our immunofluorescence results by demonstrating that expression levels of genes downstream of PERK were largely unchanged in the 5mo Tg animals compared to NTg controls and were unmodified by 414 (Fig. 3L). We measured other mediators of the UPR distinct from the PERK such as ATF6, IRE1, and other eIF2α kinases. Only the transcriptional regulator ATF6 exhibited a significant change in transcript levels in Tg mice, with a reduction of ATF6 mRNA levels in Tg + Veh mice and 414 treated mice (Supplemental Fig. 2). PERK reportedly switches preferential substrate phosphorylation in tauopathies between eIF2α and NRF2, a transcription that promotes expression of redox response proteins (*28*). 414 treatment did not alter NRF2 nuclear translocation in Tg model mice (Fig. 3M-N). Considering the off-target effects of kinase inhibitors, and those targeted by 414 beyond PERK at nanomolar and low micromolar concentrations (Table S1), we reasoned that the neuroprotective effect of 414 is likely due to a multi-target response distinct from inhibiting PERK kinase activity.

### Quantitative hippocampal proteomics reveals pathways responsible for cognitive decline in early stage tauopathy mice

To assess the effects of a compound impacting multiple cellular pathways simultaneously, we turned to quantitative proteomics to identify proteins and pathways impacted by transgenicity and our test agent, 414. Despite the prevalence of the rTg4510 model in the study of neurodegeneration, no quantitative analysis of the brain proteome in these mice has been reported. To this end, we evaluated the hippocampal proteome of NTg and Tg mice treated with either Veh or 414 (n = 4 per group, all female) using a multiplexed tandem mass tagging (TMT) approach (Fig. 4A). Following protein quantification, batch normalization, and statistical analyses, 337 proteins were identified as significantly altered across all group comparisons (FDR adjusted p value < 0.05, Fig. 4B). First, we identified the impact of 414 treatment on proteins in NTg animals (termed: “Drug Effect”) and removed these proteins from further analysis. Next, we determined which proteins were altered due to Tg genotype (termed: “Tg Effect”) by comparing NTg + Veh vs Tg + Veh samples and specifically excluded any genes known to be dysregulated due to transgene insertion in this model (*29, 30*). We then identified the proteins that were rescued by 414 treatment in the Tg mice: proteins in the “Tg Effect” group that were restored to statistically normal levels in Tg mice (no significant difference from NTg + Veh mice) following treatment with 414 (termed: “Drug Rescued”). These groupings are summarized in (Fig. 4C) and representative proteins belonging to each group are shown in (Fig. 4D). Interestingly, ≈67% of the proteins altered in Tg mice were rescued after treatment with 414 (186 out of 276). Importantly and as expected, the human *MAPT* P301L transgene was significantly upregulated in Tg mice and unaffected by 414 (Fig. 4E). Tau phosphorylation was further assessed using immobilized metal affinity chromatography (IMAC) enrichment and subsequent LC-MS/MS for phospho-proteomic analysis. Phospho-proteomic analysis revealed tau phosphorylation at 13 phospho-peptides uniquely mapping to human MAPT P301L (FDR < 1%). Averaging the abundance of these phospho-peptides as a measure of overall tau phosphorylation indicated no significant difference in tau phosphorylation in between Tg 414 treated or untreated groups (Fig. 4F). Together, these proteomic data strengthen the claim that the neuroprotective benefits of 414 are independent of tau.

**Fig 4.**
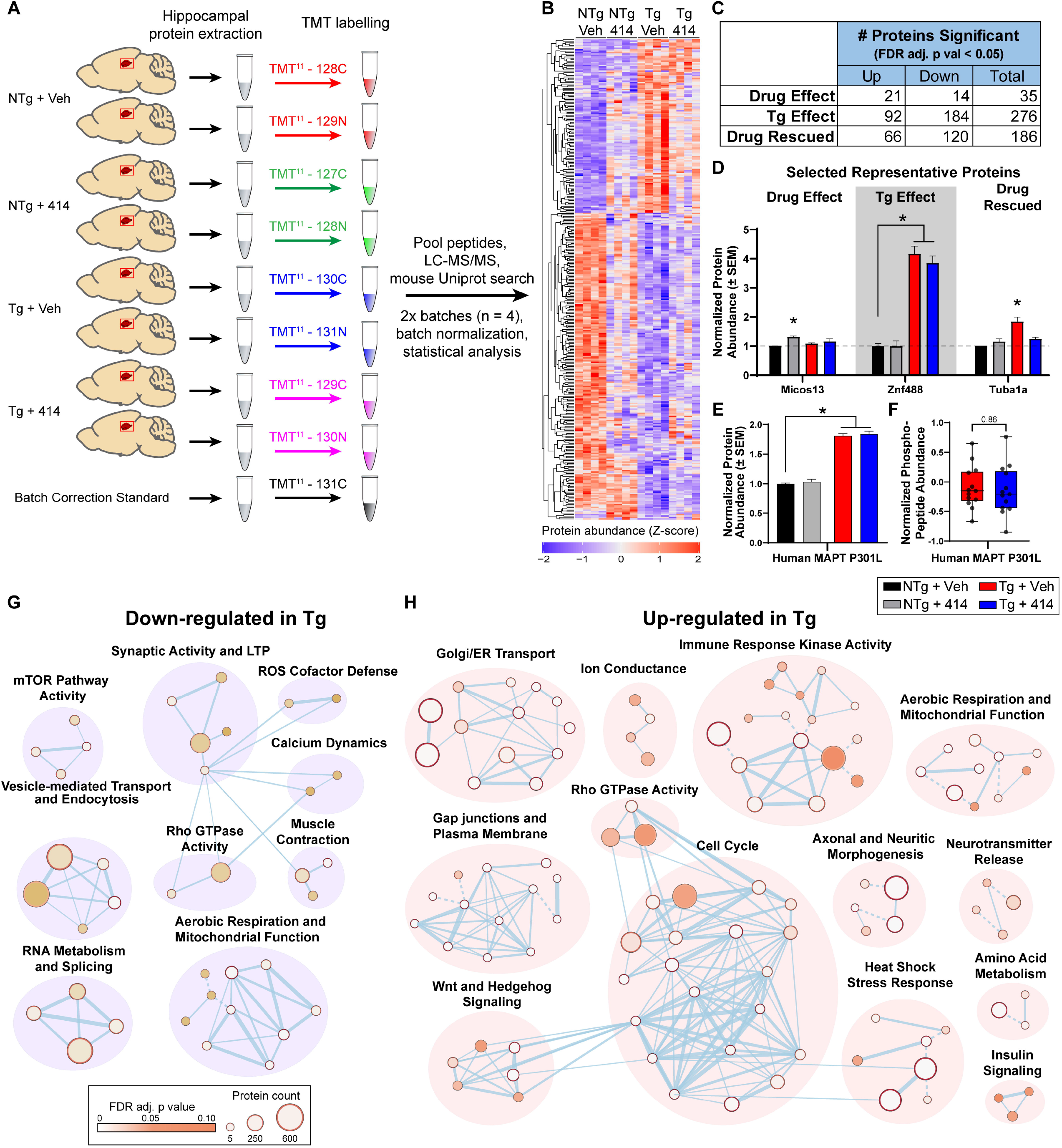
Hippocampal proteome signatures of early stage tauopathy model mice. **A** Schematic of TMT isobaric tag proteomic pipeline. Mice hippocampal sections were extracted and lysed. Individual isobaric tags were used to chemically label proteins from each individual sample and combined for LC/MS-MS detection. Batch normalization was completed using a shared batch correction standard, resulting in n = 4 per group. All samples were from female mice. **B** Heatmap of proteomics results showing 337 significantly altered proteins across all comparisons (FDR adjusted, p < 0.05). **C** Summation of the number of significantly altered proteins across each test. **D** Representative protein for each test of significant differential abundance. **E** Protein abundance of human MAPT P301L transgene. **F** Abundance of each human MAPT P301L phospho-peptide detected by phospho-proteomics. **G-H** Network clustering of Reactome pathways found up-regulated (**G**) or down-regulated (**H**) due to tau transgenic genotype relative to NTg mice (Tg effect). Node size proportionally represents the protein count in that pathway, whereas node color represents the statistical significance gradient (FDR-adjusted p value) from p = 0.10 (brown) approaching p = 0 (white). Dashed connections indicate manually adjusted nodes.

Next, we analyzed the pathway level changes in the 5mo Tg brain proteome. To collapse the altered protein into pathways, we used g.Profiler (*31*) to assess the changes in Reactome (*32*) annotated pathways (FDR adjusted p value < 0.1, Table S2). Cytoscape (*33*) was used to cluster each pathway into larger umbrella networks that were down-regulated (Fig. 4G) or up-regulated (Fig. 4H) in Tg mice. The down-regulated protein network clustered into nine discrete groups, namely: mTOR pathway activity, synaptic activity and long-term potentiation, reactive oxygen species defense, calcium dynamics, vesicle-mediated transport and endocytosis, rho GTPase activity, muscle contraction, RNA metabolism and splicing, as well as aerobic respiration and mitochondrial activity. The number of up-regulated protein networks was considerably larger despite having fewer significantly altered proteins compared to down-regulated proteins. Up-regulated pathways clustered into the following groups: Golgi/ER transport, ion conductance, immune response kinase activity, aerobic respiration and mitochondrial function, axonal and neuritic morphogenesis, neurotransmitter release, amino acid metabolism, heat shock stress response, insulin signaling, rho GTPase activity, cell cycle, Wnt and Hedgehog signaling, as well as gap junctions and plasma membrane.

### Multi-target kinase inhibition via GSK2606414 treatment rescues proteomic shifts in tauopathy

Having identified and mapped the pathway changes in Tg mice, we sought to determine the pathways that were affected by 414 that associated with cognitive rescue. To that end, we repeated clustering analysis of Reactome pathways (FDR adjusted p-value < 0.1) representing “Drug Rescued” proteins that were either initially down-regulated (Fig. 5A) or up-regulated (Fig. 5B) in Tg + Veh mice and rescued back toward NTg control levels with 414 treatment. Pathway groups initially down-regulated in Tg + Veh mice and rescued up toward NTg control levels included: aerobic respiration, long-term potentiation, muscle contraction, ion conductance, and rho GTPase activity. Other pathways initially up-regulated in Tg + Veh mice and restored to NTg control levels included: amino acid metabolism, aerobic respiration and mitochondrial function, heat shock stress response, Golgi-ER membrane trafficking, axonal and neuritic morphogenesis, apoptotic protein degradation, cell cycle, gap junction and plasma membrane, immune response kinase activity, ion conductance, NMDA post-synaptic activity, Wnt and Hedgehog signaling, and RNA metabolism and translation.

**Fig 5.**
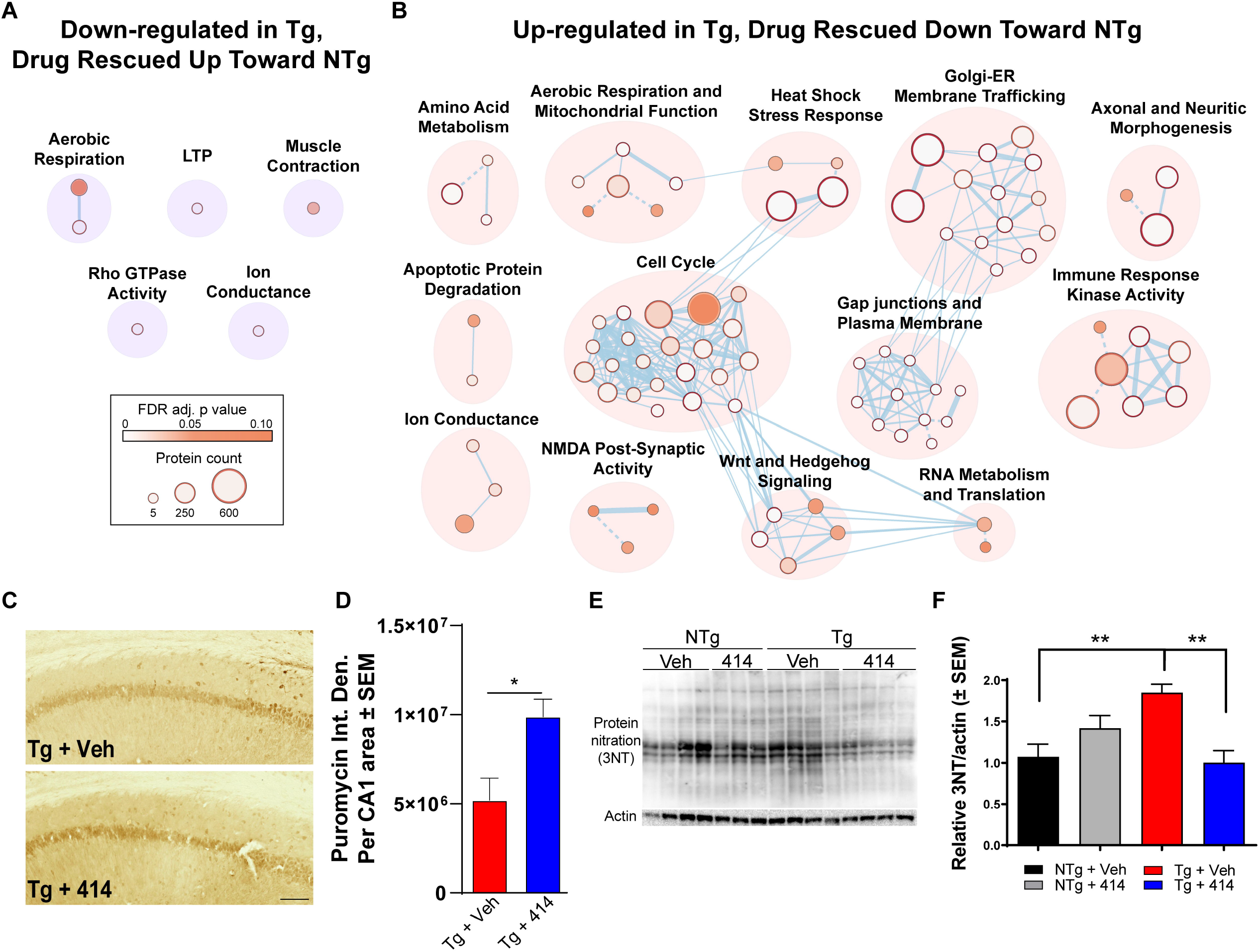
Multi-target kinase inhibition rescues pathways related to cognitive and neuronal function in early-stage tau transgenic mice. **A-B** Network clustering of Reactome pathways found up-regulated (**F**) or down-regulated (**G**) in Tg mice and significantly rescued toward NTg + Veh levels in Tg + 414 mice. Node size proportionally represents the protein count in that pathway, whereas node color represents the statistical significance gradient (FDR-adjusted p value) from p = 0.10 (brown) approaching p = 0 (white). Dashed connections indicate manually adjusted nodes. **C-D** Puromycin immunostaining of the CA1 region (**D**) and quantified (**E**) relative to total CA1 area (n = 3). One-way ANOVA with Tukey *post hoc* test. Data are expressed as the mean absolute integrated density ± SEM, *p < 0.05. Scale bar = 25 microns. **E-F** Immunoblots (e) of total tyrosine nitration and total lane quantified results (**F**) relative to NTg + Veh and normalized to actin (n = 3 - 4). Two-way ANOVA with Tukey post hoc test. Data are expressed as the mean relative to Tg + Veh ± SEM, **p < 0.01.

To validate these proteomic results, we focused on two distinct cellular processes that were rescued by 414: RNA translation and mitochondrial redox defense. RNA translation was assessed by a novel metabolic labeling method developed to investigate the translational dysregulation of tauopathy *in vivo* (*27*). This method involved an intraperitoneal injection of puromycin, a tRNA analog, shortly before death to rapidly label nascent polypeptide chains (*34*). Anti-puromycin antibodies were then used to probe for proteins that incorporated the puromycin as a direct measure of polypeptide elongation and RNA translation. Anti-puromycin staining of the CA1 region revealed significantly increased translation activity in Tg + 414 (*p < 0.05) compared to Tg + Veh mice (Fig. 5C-D). Secondly, since nitroxidative stress is a marker of mitochondrial dysregulation, and it is implicated in the pathogenesis of AD and other tauopathies (*35, 36*), we measured changes in protein nitration between our animal groups. Immunoblot results identified a significant increase in protein nitration (3-NT) in Tg + Veh animals which was rescued with 414 treatment (Fig. 5E-F). Together, these results validated the pathway level changes that were rescued in Tg mice by treatment with 414.

### Early stage tauopathy exhibits common protein alterations targetable by broad kinase inhibition

Finally, we investigated how the Tg hippocampal proteome compared with human tauopathy to provide a context for our findings in AD. We correlated the 276 proteins altered by “Tg Effect” with two independent proteomic analyses of the human AD brain to identify both model-specific alterations and novel candidate proteins (Table S3). The majority of significantly altered mouse proteins mapped to human homologs (>90%) in both human tauopathy datasets, as expected. First, to account for the disparity in pathology between the tauopathy Tg mouse model and AD, we assessed the consistency of proteomic alterations between Tg mouse and preclinical (or asymptomatic) AD and late-stage AD samples (Fig. 6A). Individuals with asymptomatic AD (Asym AD) included in this analysis exhibited minimal cognitive impairment with moderate to high amyloid plaque frequency, but with low to no cortical tau pathology (Table S4) (*37*). Late AD samples, however, had significant cognitive impairment and high degrees of amyloid and tau pathology. By identifying the proteins consistent between Tg mice and Asym AD distinct from late AD, we classified which proteins are considered early markers of AD- and tauopathy-specific progression, respectively. We found that 15 of 20 human homologs significantly altered in Asym AD vs healthy control brains were consistently altered in Tg animals (75%; hypergeometric test, ****p < 0.0001). Of these 15, nine homolog proteins were rescued by 414 in Tg mice: NUCB1, SNCA, FKBP1A, PKM, YWHAZ, HSPE1, GRPEL1, NDUFB10, and HOMER1. These nine proteins are implicated in other neurodegenerative diseases or cellular processes known to be dysregulated early in tauopathy pathogenesis such as synaptic plasticity, proteostasis, glucose metabolism, and mitochondrial function. Tg mice matched to a greater extent with the late AD proteome, where 29 out of 47 significantly altered proteins were inter-specially consistent (≈62%; hypergeometric test, **p < 0.01). Of these 29 proteins, 23 homologs were rescued by 414 in Tg mice: NUCB1, NTM, ATOX1, SNCA, ENO1, GJA1, FKBP1A, ATP6V1E1, PKM, STMN1, VSNL1, YWHAZ, HINT1, HPCA, PEPD, ROGDI, PPP1R7, PACSIN1, HSPE1, RTN1, NDUFB5, DMTN, and HOMER1. These proteins function in pathways rescued by 414 in Tg mice, namely aerobic respiration/glucose metabolism, mitochondrial function, endocytosis and trafficking, immune system activation, signaling cascades. Interestingly, several proteins were found consistent across disease staging: NUCB1, SNCA, FKBP1A, PKM, YWHAZ, HSPE1, HOMER1. Since each of these were rescued by 414 in Tg mice, they may be important candidate proteins to consider for future studies.

**Fig 6:**
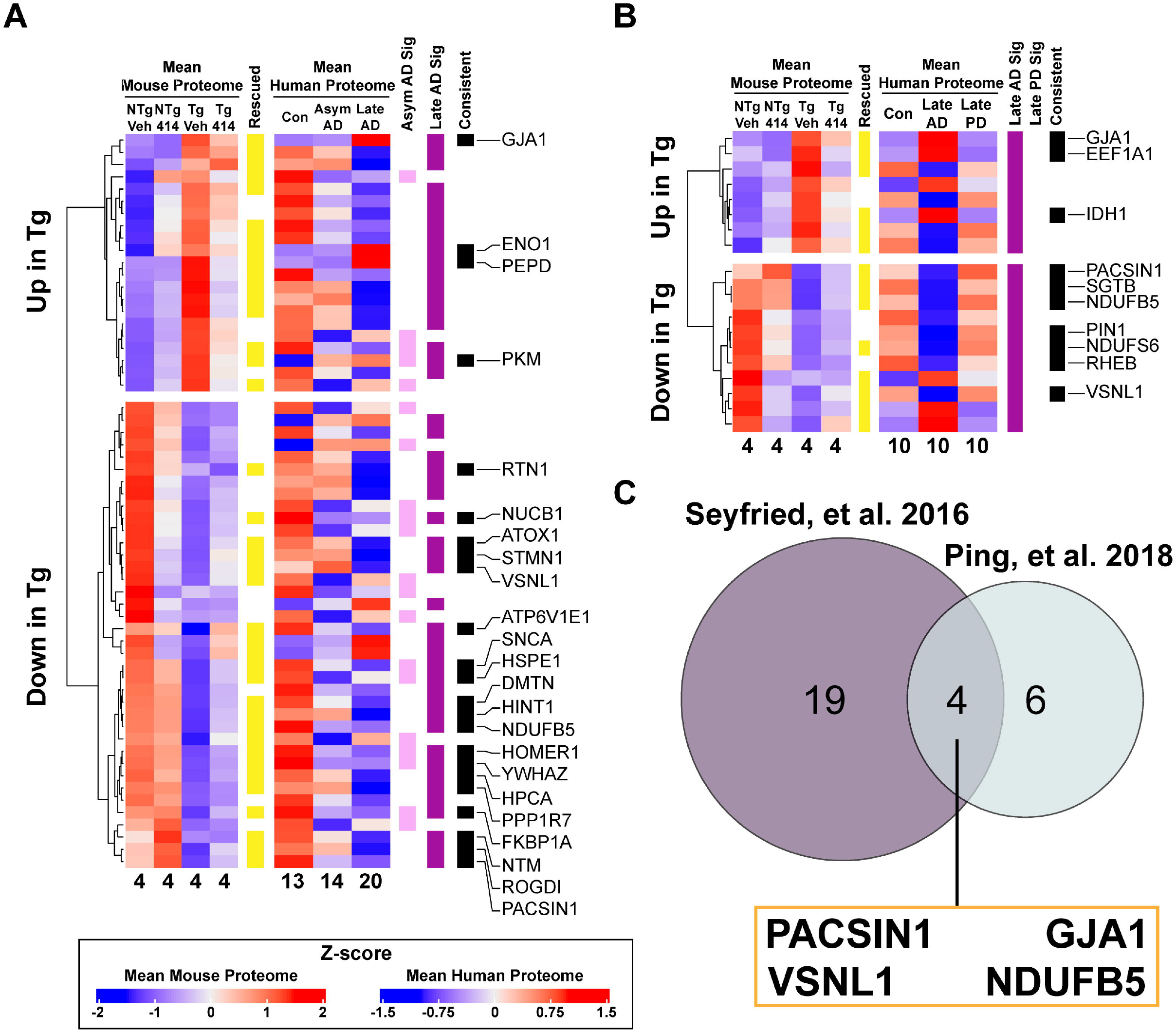
Proteomic signatures of rTg4510 transgenic mice match human tauopathy and reveal novel candidates for therapeutic intervention. **A-B** Significantly altered proteins by Tg effect were compared to human homolog abundances found in two proteomics datasets of human neurodegenerative diseases (**A**, (*37*); **B**, (*39*)). Proteins annotated in the “Rescued” column (yellow) were significantly rescued by 414 in Tg animals compared to NTg controls. Human homolog protein abundances in each disease group were compared to non-demented controls. Statistically altered human proteins were annotated by disease group in different colors (Asym AD, pink; late AD, purple; late PD, green). Proteins that were significantly altered in the same direction (increased or decreased) between species and also represented proteins rescued by 414 in Tg mice were annotated in the “Consistent” column (black). Candidate proteins passing all criteria for inclusion were labeled with their corresponding human gene symbols. Sample abundances were averaged per group and each corresponding sample number was appended to the bottom of each column. **C** Venn diagram comparing candidate proteins identified in late AD across both datasets.

Considering that the proteome in late-stage neurodegenerative diseases may relate more to widespread brain organ failure and may not be a specific response to any one disease (*38*), we investigated how the Tg proteome matched the protein alterations in AD and another chronic, neurodegenerative disease, Parkinson’s disease (PD) (*39*). We applied a similar approach used with asymptomatic and late stage AD, wherein we analyzed whether proteins found altered in the Tg hippocampal proteome were consistently altered in AD or PD (Fig. 6B). Interestingly, Tg animals and PD brains did not share consistently altered homologous proteins, suggesting a tauopathy specific effect. Of the 19 significantly altered proteins found in AD, 12 had homologs consistently changed in Tg animals (≈63%; hypergeometric test *p<0.05). Of these twelve proteins, 10 were rescued by 414: six unique to Tg animals (GJA1, EEF1A1, IDH1, SGTB, PIN1, RHEB) and four proteins that are shared across both late AD (human and Tg) datasets: PACSIN1, GJA1, VSNL1, and NDUFB5 (Fig. 6C). These high-confidence candidate proteins associated with cognitive decline and drug-mediated rescue are. Further analyses are required to address the role of these candidate genes in the

## Discussion

Here, we uncover critical biological processes that contribute to neurotoxic processes driving cognitive dysfunction in tauopathies using a broad kinase inhibitor, GSK2606414. In doing so, we demonstrate the potential to mitigate the negative functional consequences of tauopathy without altering tau hyper-phosphorylation. We treated early stage rTg4510 mice and firmly establish cognitive and molecular benefits of broad kinase inhibition. With TMT quantitative proteomics, we identified biological processes that are affected in transgenics and rescued by the compound. Finally, we cross-reference our findings to human AD and PD brain proteomic data to narrow our results to high confidence unique targets for future study. This approach revealed four distinct proteins associated with cognitive decline in human tauopathy with strong therapeutic potential.

GSK2606414 ameliorates molecular underpinnings of neurodegeneration including neuronal loss, cognitive dysfunction, and pathological protein accumulation in a variety of disease models (*11–16, 18*). These studies established that the benefits of GSK2606414 were conferred by inhibition of its primary target, PERK, which was upregulated in disease. While PERK is active in the rTg4510 tau transgenic model, this does not occur until after 6mo (*11, 26*). By treating rTg4510 mice from 4-5mo, a time point in which PERK levels not activated, our study identified numerous PERK-independent pathways targeted by GSK2606414 that elicited rescue from toxic outcomes in tauopathy. This has important implications for data interpretation of this and previous studies.

Many pathways affected by disease in this early stage of rTg4510 mice overlap with altered pathways in AD brain such as glucose metabolism, immune response, synaptic activity, the MAPK cascade, and RNA metabolism, highlighting the similarity between this model and human tauopathy (*6–8, 37*). Comparing our proteomic results with a study of proteostatic dysfunction and protein aggregation in rTg4510 reveals no apparent correlation of protein abundance between soluble and insoluble fractions (*40*). These studies suggest that our proteomic findings reflect alterations in expression and degradation rather than a protein shift toward insolubility.

Given the congruent pathway-level results between rTg4510 mice and human tauopathy studies, we searched for novel candidate targets that were associated with rescued cognitive function with GSK2606414 treatment. Overlapping the proteins consistent in both AD datasets with data from transgenic mice revealed four novel, high-confidence candidates that were associated with cognitive decline and drug-mediated multi-target kinase inhibition: PACSIN1, GJA1, VSNL1, and NDUFB5. The neuronal protein PACSIN1, or protein kinase C and casein kinase substrate in neurons protein 1, participates in endocytosis (*41–43*) and directly interacts with tubulin to promote microtubule formation (*44*). There is evidence for direct tau-PACSIN1 binding, suggesting that this interaction impinges upon tau’s capacity to bind microtubules (*45*) and represents a protein with a direct link to tauopathy that could perturb cellular trafficking and tau/microtubule stability. GJA1/connexin43, or gap junction alpha 1, functions to allow rapid inter-cellular communication of small molecules such as ions and neurotransmitters by establishing hemichannel gap junctions between cells (*46*). Many studies coupled GJA1 with the pathogenesis of neurodegenerative diseases such as AD (*47, 48*), most notably finding GJA1 incorporated into amyloid plaques (*49*) and upregulated in AD models and human brain (*6, 50– 52*). VSNL1, or visinin-like protein 1, is a neuronal calcium sensor (NCS) protein highly expressed in the brain. As an NCS, VSNL1 responds to alterations in calcium concentration and coordinates physiological processes such as cyclic nucleotide second messenger cascades and synaptic receptor recycling (*53–55*). While the full functional role of VSNL1 remains unclear, multiple studies have implicated VSNL1 in progressing calcium and synaptic dysfunction in AD (*56*), stroke (*57*), and schizophrenia (*58*). Biomarker studies found increased VSNL1 in CSF and plasma in early AD which correlated with cognitive dysfunction and neuronal loss (*59–62*). NDUFB5, or nicotinamide adenine dinucleotide dehydrogenase (ubiquinone) 1 beta subcomplex 5, is a nuclear-encoded mitochondrial protein which functions as part of complex 1 of the electron transport chain. Many studies have shown NDUFB5 levels is altered in disease, typically coupled with altered expression of other mitochondrial respiratory proteins (*63–66*). In the context of AD, NDUFB5 has been suggested as a hub protein associated with disease pathogenesis based on co-expression analyses (*67, 68*).

Taken together, our pre-clinical study contributes two major findings. First, we identify discrete pathways contributing to cognitive changes in tauopathy and highlight four unique proteins associated with cognitive rescue with strong therapeutic potential. Further work investigating the novel targets identified in this study could discover better targets amenable to pharmacological intervention and novel involvement in tauopathy. Second, we demonstrate that broad, multi-family kinase inhibition can be a useful tool to mitigate the molecular and functional decline in tauopathy. Since extensively multi-targeted approaches meet safety challenges in the clinic, the development of compounds that selectively target key kinase cascades at the same time could greatly benefit combinatorial therapies for the treatment of tauopathy. Studies combining FDA-approved selective kinase inhibitors may be beneficial toward advancing possible therapeutic strategies.

## Materials and Methods

### Study Design

The objective of this study was to determine the efficacy of broadly inhibiting kinase activity for the treatment of early stage tauopathy. The rationale is that kinases are implicated in the early pathogenesis and progression of tauopathies, such as AD, though our understanding of the dysregulated kinase network in these diseases originates from studies isolating the effect of one kinase or kinase family. Given the multi-factorial complexity of these diseases, we investigated whether the considerable off-target effects of a pharmacological agent (*69*) could provide clinical benefit rather than act to confound experiments. Therefore, we evaluated the effect of a compound originally developed as a selective inhibitor for PERK (*10*) and used in that context in a variety of disease models (*11, 12, 16*), but which reportedly broadly targets kinases at low micromolar concentrations (*10, 17, 18*). We tested the outcomes of treating tau transgenic mice with this compound across a variety of functional, cognitive, and molecular measures of tauopathy. We confirmed that the neuroprotective effect of the compound is independent of PERK activity at this early age, though older mice with PERK activity respond to the compound as previously reported. Lastly, we investigated the protein changes underlying the cognitive dysfunction and subsequent rescue at this age in tau transgenic mice. To control for transgene-specific effects in mice which may not be recapitulated tauopathy in humans, we compared these results with two separate measures of the AD proteome using different proteomic techniques to control for potential systematic biases.

All experiments were designed with appropriate controls and based on previous experiments to determine statistical power (*22, 27, 70*). Animals were separated by genotype and randomly assigned to treatment groups and experimental cohorts. Mice which did not undergo full treatment course, such as those removed from analysis due to body weight reduction past 80% of starting weight, were not analyzed for this study.

### Animals

All animal studies were approved by the University of Kentucky’s Institutional Animal Care and Use Committee (IACUC) and abided by that committee’s policies on animal care and use in accordance with federal guidelines. Mice were kept in standard housing on a 12h light/dark cycle and received food and water ad libitum. The tau transgenic (rTg4510) and parental mice were maintained and genotyped as described previously (*20, 27*) and were maintained on mixed FVB and 129S6 backgrounds. Mice of both sexes were used in experiments unless otherwise stated.

### Treatment with GSK2606414

GSK2606414 (GlaxoSmithKline) was suspended in vehicle (0.5% hydroxypropylmethyl cellulose + 0.1% Tween-80 in water at pH 4.0) as previously described (*12*). A total of 100 mg/kg GSK2606414 was delivered by oral gavage once (100 mg/kg) or twice a day (50 mg/kg each separated by 10-12 hours). Prior to treatment, mice were handled twice daily to acclimate the animals for gavage. Unless otherwise stated, four-month-old rTg4510 and non-transgenic mice were treated with GSK2606414 or vehicle for 30-33 days ranging to 36 days to accommodate for environmental habituation for behavioral and MRI assays. Daily weight records were kept to ensure accurate dosing and to monitor potential GSK2606414-mediated weight loss (*12*). Animals that lost more than 20% of starting body weight were excluded from the study (Supplemental Fig. 3). No group exhibited pronounced weight loss.

### Statistical Analysis

Statistical analyses for all data apart from proteomics were performed using GraphPad Prism 8 (Graph Pad Software, Inc. La Jolla, CA, USA). Results are shown as the mean ± standard error or standard deviation as described in each figure. Single-variate data were analyzed using unpaired Student’s t-test. Multi-variate data were analyzed with one-way or two-way ANOVA where appropriate, corrected for multiple comparisons with Tukey post-test analysis unless otherwise stated. A value of p < 0.05 was considered statistically significant except for Reactome pathway analyses which used an FDR-adjusted p value cutoff of 0.10.

## Acknowledgements

We thank the UF Proteomics and Mass Spectrometry core for proteomic analysis, the University of Kentucky Magnetic Resonance Imaging and Spectroscopy Center (MRISC) for MRI analysis. We thank Dr. Peter Davies for his generous contribution of the PHF1 antibody. We thank Dr. Fred Schmitt for insightful discussions and crucial intellectual contributions to this study. We thank Dr. Shelby Meier for her contributions to the work prior to this study and experimental support in this current project.

## Author Contributions

S.A.K., S.N.F., and J.F.A. designed the project. S.A.K., M.J.H., R.C., S.N.F., C.L., J.R.R., A.I., M.B., E.J.M., S.M.G.E., N.Z., N.S., D.T., D.P., M.V., T.F., and A.M.S.H. performed experiments, acquired data, or assisted in analysis. S.A.K., M.J.H., and J.F.A drafted the manuscript. J.K., J.M.A., and N.J.L. edited the manuscript.

## Data and materials availability

Raw proteomic data will be accessible in PRIDE.

## Supplementary Materials

### Supplementary Materials and Methods

Fig. S1. Total duration spent in novel arm is partially rescued in tau transgenic mice treated with GSK2606414.

Fig. S2. Non-PERK UPR proteins do not have altered transcript levels in 5mo tau transgenic mice.

Fig. S3. GSK2606414 treatment for up to 36d does not cause marked weight loss.

Table S1. Top kinase targets of GSK2606414 and relevance to tauopathy.

Table S2. Significantly enriched reactome pathways identified by proteomics.

Table S3. Human to mouse proteomic comparisons.

Table S4. Patient demographics of human proteomic samples.

References (*71, 72*)

## References

1. M. R. Khanna, J. Kovalevich, V. M.-Y. Lee, J. Q. Trojanowski, K. R. Brunden, Therapeutic Strategies for the Treatment of Tauopathies: Hopes and Challenges, Alzheimers Dement 12, 1051–1065 (2016).

2. S. Jadhav, J. Avila, M. Schöll, G. G. Kovacs, E. Kövari, R. Skrabana, L. D. Evans, E. Kontsekova, B. Malawska, R. de Silva, L. Buee, N. Zilka, A walk through tau therapeutic strategies, Acta Neuropathologica Communications 7, 22 (2019).

3. L.-K. Huang, S.-P. Chao, C.-J. Hu, Clinical trials of new drugs for Alzheimer disease, Journal of Biomedical Science 27, 18 (2020).

4. A. L. Boxer, A. E. Lang, M. Grossman, D. S. Knopman, B. L. Miller, L. S. Schneider, R. S. Doody, Lees, L. I. Golbe, D. R. Williams, J.-C. Corvol, A. Ludolph, D. Burn, S. Lorenzl, I. Litvan, E. D. Roberson, G. U. Höglinger, M. Koestler, C. R. Jack, V. Van Deerlin, C. Randolph, I. V. Lobach, H. W. Heuer, I. Gozes, L. Parker, S. Whitaker, J. Hirman, A. J. Stewart, M. Gold, B. H. Morimoto, AL-108-231 Investigators, Davunetide in patients with progressive supranuclear palsy: a randomised, double-blind, placebo-controlled phase 2/3 trial, Lancet Neurol 13, 676–685 (2014).

5. R. R. Ramsay, M. R. Popovic-Nikolic, K. Nikolic, E. Uliassi, M. L. Bolognesi, A perspective on multi-target drug discovery and design for complex diseases, Clinical and Translational Medicine 7, 3 (2018).

6. E. C. B. Johnson, E. B. Dammer, D. M. Duong, L. Ping, M. Zhou, L. Yin, L. A. Higginbotham, A. Guajardo, B. White, J. C. Troncoso, M. Thambisetty, T. J. Montine, E. B. Lee, J. Q. Trojanowski, T. G. Beach, E. M. Reiman, V. Haroutunian, M. Wang, E. Schadt, B. Zhang, D. W. Dickson, N. Ertekin-Taner, T. E. Golde, V. A. Petyuk, P. L. De Jager, D. A. Bennett, T. S. Wingo, S. Rangaraju, I. Hajjar, J. M. Shulman, J. J. Lah, A. I. Levey, N. T. Seyfried, Large-scale proteomic analysis of Alzheimer’s disease brain and cerebrospinal fluid reveals early changes in energy metabolism associated with microglia and astrocyte activation, Nat. Med. 26, 769–780 (2020).

7. E. C. B. Johnson, E. B. Dammer, D. M. Duong, L. Yin, M. Thambisetty, J. C. Troncoso, J. J. Lah, I. Levey, N. T. Seyfried, Deep proteomic network analysis of Alzheimer’s disease brain reveals alterations in RNA binding proteins and RNA splicing associated with disease, Mol Neurodegener 13, 52 (2018).

8. B. Bai, X. Wang, Y. Li, P.-C. Chen, K. Yu, K. K. Dey, J. M. Yarbro, X. Han, B. M. Lutz, S. Rao, Y. Jiao, J. M. Sifford, J. Han, M. Wang, H. Tan, T. I. Shaw, J.-H. Cho, S. Zhou, H. Wang, M. Niu, A. Mancieri, K. A. Messler, X. Sun, Z. Wu, V. Pagala, A. A. High, W. Bi, H. Zhang, H. Chi, V. Haroutunian, B. Zhang, T. G. Beach, G. Yu, J. Peng, Deep Multilayer Brain Proteomics Identifies Molecular Networks in Alzheimer’s Disease Progression, Neuron 105, 975-991.e7 (2020).

9. M. Perluigi, E. Barone, F. Di Domenico, D. A. Butterfield, Aberrant protein phosphorylation in Alzheimer disease brain disturbs pro-survival and cell death pathways, Biochim. Biophys. Acta 1862, 1871–1882 (2016).

10. J. M. Axten, J. R. Medina, Y. Feng, A. Shu, S. P. Romeril, S. W. Grant, W. H. H. Li, D. A. Heerding, E. Minthorn, T. Mencken, C. Atkins, Q. Liu, S. Rabindran, R. Kumar, X. Hong, A. Goetz, T. Stanley, J. D. Taylor, S. D. Sigethy, G. H. Tomberlin, A. M. Hassell, K. M. Kahler, L. M. Shewchuk, R. T. Gampe, Discovery of 7-methyl-5-(1-{[3-(trifluoromethyl)phenyl]acetyl}-2,3-dihydro-1H-indol-5-yl)-7H-pyrrolo[2,3-d]pyrimidin-4-amine (GSK2606414), a potent and selective first-in-class inhibitor of protein kinase R (PKR)-like endoplasmic reticulum kinase (PERK), J. Med. Chem. 55, 7193–7207 (2012).

11. H. Radford, J. A. Moreno, N. Verity, M. Halliday, G. R. Mallucci, PERK inhibition prevents tau-mediated neurodegeneration in a mouse model of frontotemporal dementia, Acta Neuropathol 130, 633–642 (2015).

12. J. A. Moreno, M. Halliday, C. Molloy, H. Radford, N. Verity, J. M. Axten, C. A. Ortori, A. E. Willis, P. M. Fischer, D. A. Barrett, G. R. Mallucci, Oral treatment targeting the unfolded protein response prevents neurodegeneration and clinical disease in prion-infected mice, Sci Transl Med 5, 206ra138 (2013).

13. C. Meng, J. Zhang, B. Dang, H. Li, H. Shen, X. Li, Z. Wang, PERK Pathway Activation Promotes Intracerebral Hemorrhage Induced Secondary Brain Injury by Inducing Neuronal Apoptosis Both in Vivo and in Vitro, Front Neurosci 12 (2018), doi: 10.3389/fnins.2018.00111.

14. T. Vanderweyde, D. J. Apicco, K. Youmans-Kidder, P. E. A. Ash, C. Cook, E. L. da Rocha, K. Jansen-West, A. A. Frame, A. Citro, J. D. Leszyk, P. Ivanov, J. F. Abisambra, M. Steffen, H. Li, L. Petrucelli, B. Wolozin, Interaction of tau with the RNA-Binding Protein TIA1 Regulates tau Pathophysiology and Toxicity, Cell Rep 15, 1455–1466 (2016).

15. V. Sharma, H. Ounallah-Saad, D. Chakraborty, M. Hleihil, R. Sood, I. Barrera, E. Edry, S. K. Chandran, S. B. T. de Leon, H. Kaphzan, K. Rosenblum, Local Inhibition of PERK Enhances Memory and Reverses Age-Related Deterioration of Cognitive and Neuronal Properties, J. Neurosci. 38, 648–658 (2018).

16. H. L. Smith, O. J. Freeman, A. J. Butcher, S. Holmqvist, I. Humoud, T. Schätzl, D. T. Hughes, N. C. Verity, D. P. Swinden, J. Hayes, L. de Weerd, D. H. Rowitch, R. J. M. Franklin, G. R. Mallucci, Astrocyte Unfolded Protein Response Induces a Specific Reactivity State that Causes Non-Cell-Autonomous Neuronal Degeneration, Neuron 105, 855-866.e5 (2020).

17. D. Rojas-Rivera, T. Delvaeye, R. Roelandt, W. Nerinckx, K. Augustyns, P. Vandenabeele, M. J. M. Bertrand, When PERK inhibitors turn out to be new potent RIPK1 inhibitors: critical issues on the specificity and use of GSK2606414 and GSK2656157, Cell Death Differ 24, 1100–1110 (2017).

18. M. Mahameed, T. Wilhelm, O. Darawshi, A. Obiedat, W.-S. Tommy, C. Chintha, T. Schubert, A. Samali, E. Chevet, L. A. Eriksson, M. Huber, B. Tirosh, The unfolded protein response modulators GSK2606414 and KIRA6 are potent KIT inhibitors, Cell Death Dis 10, 300 (2019).

19. Z. Berger, H. Roder, A. Hanna, A. Carlson, V. Rangachari, M. Yue, Z. Wszolek, K. Ashe, J. Knight, D. Dickson, C. Andorfer, T. L. Rosenberry, J. Lewis, M. Hutton, C. Janus, Accumulation of Pathological Tau Species and Memory Loss in a Conditional Model of Tauopathy, J. Neurosci. 27, 3650–3662 (2007).

20. K. SantaCruz, J. Lewis, T. Spires, J. Paulson, L. Kotilinek, M. Ingelsson, A. Guimaraes, M. DeTure, M. Ramsden, E. McGowan, C. Forster, M. Yue, J. Orne, C. Janus, A. Mariash, M. Kuskowski, B. Hyman, M. Hutton, K. H. Ashe, Tau Suppression in a Neurodegenerative Mouse Model Improves Memory Function, Science 309, 476–481 (2005).

21. R. M. Bailey, J. Howard, J. Knight, N. Sahara, D. W. Dickson, J. Lewis, Effects of the C57BL/6 strain background on tauopathy progression in the rTg4510 mouse model, Molecular Neurodegeneration 9, 8 (2014).

22. S. N. Fontaine, A. Ingram, R. A. Cloyd, S. E. Meier, E. Miller, D. Lyons, G. K. Nation, E. Mechas, B. Weiss, C. Lanzillotta, F. Di Domenico, F. Schmitt, D. K. Powell, M. Vandsburger, J. F. Abisambra, Identification of changes in neuronal function as a consequence of aging and tauopathic neurodegeneration using a novel and sensitive magnetic resonance imaging approach, Neurobiol Aging 56, 78–86 (2017).

23. F. M. LaFerla, Calcium dyshomeostasis and intracellular signalling in Alzheimer’s disease, Nat. Rev. Neurosci. 3, 862–872 (2002).

24. C. Hooper, R. Killick, S. Lovestone, The GSK3 hypothesis of Alzheimer’s disease, J Neurochem 104, 1433–1439 (2008).

25. M. C. Bell, S. E. Meier, A. L. Ingram, J. F. Abisambra, PERK-opathies: An Endoplasmic Reticulum Stress Mechanism Underlying Neurodegeneration, Curr Alzheimer Res 13, 150–163 (2016).

26. J. F. Abisambra, U. K. Jinwal, L. J. Blair, J. C. O’Leary, Q. Li, S. Brady, L. Wang, C. E. Guidi, B. Zhang, B. A. Nordhues, M. Cockman, A. Suntharalingham, P. Li, Y. Jin, C. A. Atkins, C. A. Dickey, Tau accumulation activates the unfolded protein response by impairing endoplasmic reticulum-associated degradation, J. Neurosci. 33, 9498–9507 (2013).

27. S. A. Koren, M. J. Hamm, S. E. Meier, B. E. Weiss, G. K. Nation, E. A. Chishti, J. P. Arango, J. Chen, H. Zhu, E. M. Blalock, J. F. Abisambra, Tau drives translational selectivity by interacting with ribosomal proteins, Acta Neuropathol 137, 571–583 (2019).

28. S. B. Cullinan, D. Zhang, M. Hannink, E. Arvisais, R. J. Kaufman, J. A. Diehl, Nrf2 is a direct PERK substrate and effector of PERK-dependent cell survival, Mol. Cell. Biol. 23, 7198–7209 (2003).

29. L. O. Goodwin, E. Splinter, T. L. Davis, R. Urban, H. He, R. E. Braun, E. J. Chesler, V. Kumar, M. van Min, J. Ndukum, V. M. Philip, L. G. Reinholdt, K. Svenson, J. K. White, M. Sasner, C. Lutz, S. A. Murray, Large-scale discovery of mouse transgenic integration sites reveals frequent structural variation and insertional mutagenesis, Genome Res. 29, 494–505 (2019).

30. J. Gamache, K. Benzow, C. Forster, L. Kemper, C. Hlynialuk, E. Furrow, K. H. Ashe, M. D. Koob, Factors other than hTau overexpression that contribute to tauopathy-like phenotype in rTg4510 mice, Nat Commun 10, 2479 (2019).

31. U. Raudvere, L. Kolberg, I. Kuzmin, T. Arak, P. Adler, H. Peterson, J. Vilo, g:Profiler: a web server for functional enrichment analysis and conversions of gene lists (2019 update), Nucleic Acids Res 47, W191–W198 (2019).

32. B. Jassal, L. Matthews, G. Viteri, C. Gong, P. Lorente, A. Fabregat, K. Sidiropoulos, J. Cook, M. Gillespie, R. Haw, F. Loney, B. May, M. Milacic, K. Rothfels, C. Sevilla, V. Shamovsky, S. Shorser, T. Varusai, J. Weiser, G. Wu, L. Stein, H. Hermjakob, P. D’Eustachio, The reactome pathway knowledgebase, Nucleic Acids Res. 48, D498–D503 (2020).

33. P. Shannon, A. Markiel, O. Ozier, N. S. Baliga, J. T. Wang, D. Ramage, N. Amin, B. Schwikowski, T. Ideker, Cytoscape: a software environment for integrated models of biomolecular interaction networks, Genome Res. 13, 2498–2504 (2003).

34. C. A. Goodman, T. A. Hornberger, Measuring protein synthesis with SUnSET: a valid alternative to traditional techniques?, Exerc Sport Sci Rev 41, 107–115 (2013).

35. D. A. Butterfield, T. T. Reed, M. Perluigi, C. De Marco, R. Coccia, J. N. Keller, W. R. Markesbery, R. Sultana, Elevated Levels of 3-Nitrotyrosine in Brain From Subjects with Amnestic Mild Cognitive Impairment: Implications for the Role of Nitration in the Progression of Alzheimer’s Disease, Brain Res 1148, 243–248 (2007).

36. D. A. Butterfield, T. Reed, R. Sultana, Roles of 3-nitrotyrosine- and 4-hydroxynonenal-modified brain proteins in the progression and pathogenesis of Alzheimer’s disease, Free Radic. Res. 45, 59–72 (2011).

37. N. T. Seyfried, E. B. Dammer, V. Swarup, D. Nandakumar, D. M. Duong, L. Yin, Q. Deng, T. Nguyen, C. M. Hales, T. Wingo, J. Glass, M. Gearing, M. Thambisetty, J. C. Troncoso, D. H. Geschwind, J. J. Lah, A. I. Levey, A Multi-network Approach Identifies Protein-Specific Co-expression in Asymptomatic and Symptomatic Alzheimer’s Disease, Cell Syst 4, 60-72.e4 (2017).

38. T. E. Golde, S. T. DeKosky, D. Galasko, Alzheimer’s disease: The right drug, the right time, Science 362, 1250–1251 (2018).

39. L. Ping, D. M. Duong, L. Yin, M. Gearing, J. J. Lah, A. I. Levey, N. T. Seyfried, Global quantitative analysis of the human brain proteome in Alzheimer’s and Parkinson’s Disease, Sci Data 5, 180036 (2018).

40. M. C. Pace, G. Xu, S. Fromholt, J. Howard, K. Crosby, B. I. Giasson, J. Lewis, D. R. Borchelt, Changes in proteome solubility indicate widespread proteostatic disruption in mouse models of neurodegenerative disease, Acta Neuropathol 136, 919–938 (2018).

41. J. Modregger, B. Ritter, B. Witter, M. Paulsson, M. Plomann, All three PACSIN isoforms bind to endocytic proteins and inhibit endocytosis, J. Cell. Sci. 113 Pt 24, 4511–4521 (2000).

42. V. Anggono, K. J. Smillie, M. E. Graham, V. A. Valova, M. A. Cousin, P. J. Robinson, Syndapin I is the phosphorylation-regulated dynamin I partner in synaptic vesicle endocytosis, Nat. Neurosci. 9, 752–760 (2006).

43. A. Braun, R. Pinyol, R. Dahlhaus, D. Koch, P. Fonarev, B. D. Grant, M. M. Kessels, B. Qualmann, EHD proteins associate with syndapin I and II and such interactions play a crucial role in endosomal recycling, Mol. Biol. Cell 16, 3642–3658 (2005).

44. E.-M. S. Grimm-Günter, M. Milbrandt, B. Merkl, M. Paulsson, M. Plomann, PACSIN proteins bind tubulin and promote microtubule assembly, Exp. Cell Res. 314, 1991–2003 (2008).

45. Y. Liu, K. Lv, Z. Li, A. C. H. Yu, J. Chen, J. Teng, PACSIN1, a Tau-interacting protein, regulates axonal elongation and branching by facilitating microtubule instability, J. Biol. Chem. 287, 39911–39924 (2012).

46. D. A. Goodenough, D. L. Paul, Beyond the gap: functions of unpaired connexon channels, Nat. Rev. Mol. Cell Biol. 4, 285–294 (2003).

47. J.-T. He, X.-Y. Li, L. Yang, X. Zhao, Astroglial connexins and cognition: memory formation or deterioration?, Biosci Rep 40 (2020), doi: 10.1042/BSR20193510.

48. L. C. Mayorquin, A. V. Rodriguez, J.-J. Sutachan, S. L. Albarracín, Connexin-Mediated Functional and Metabolic Coupling Between Astrocytes and Neurons, Front Mol Neurosci 11 (2018), doi: 10.3389/fnmol.2018.00118.

49. J. I. Nagy, W. Li, E. L. Hertzberg, C. A. Marotta, Elevated connexin43 immunoreactivity at sites of amyloid plaques in Alzheimer’s disease, Brain Res. 717, 173–178 (1996).

50. R. Ren, L. Zhang, M. Wang, Specific deletion connexin43 in astrocyte ameliorates cognitive dysfunction in APP/PS1 mice, Life Sci. 208, 175–191 (2018).

51. X. Mei, P. Ezan, C. Giaume, A. Koulakoff, Astroglial connexin immunoreactivity is specifically altered at β-amyloid plaques in β-amyloid precursor protein/presenilin1 mice, Neuroscience 171, 92–105 (2010).

52. Y. Kajiwara, E. Wang, M. Wang, W. C. Sin, K. J. Brennand, E. Schadt, C. C. Naus, J. Buxbaum, B. Zhang, GJA1 (connexin43) is a key regulator of Alzheimer’s disease pathogenesis, Acta Neuropathol Commun 6, 144 (2018).

53. M. Brackmann, S. Schuchmann, R. Anand, K.-H. Braunewell, Neuronal Ca2+ sensor protein VILIP-1 affects cGMP signalling of guanylyl cyclase B by regulating clathrin-dependent receptor recycling in hippocampal neurons, J. Cell. Sci. 118, 2495–2505 (2005).

54. R. D. Burgoyne, Neuronal calcium sensor proteins: generating diversity in neuronal Ca2+ signalling, Nat. Rev. Neurosci. 8, 182–193 (2007).

55. K.-H. Braunewell, A. J. Klein-Szanto, Visinin-like proteins (VSNLs): interaction partners and emerging functions in signal transduction of a subfamily of neuronal Ca2+-sensor proteins, Cell Tissue Res 335, 301–316 (2009).

56. M. Groblewska, P. Muszyński, A. Wojtulewska-Supron, A. Kulczyńska-Przybik, B. Mroczko, The Role of Visinin-Like Protein-1 in the Pathophysiology of Alzheimer’s Disease, J. Alzheimers Dis. 47, 17–32 (2015).

57. O. F. Laterza, V. R. Modur, D. L. Crimmins, J. V. Olander, Y. Landt, J.-M. Lee, J. H. Ladenson, Identification of novel brain biomarkers, Clin. Chem. 52, 1713–1721 (2006).

58. K. H. Braunewell, A. D. Dwary, F. Richter, K. Trappe, C. Zhao, I. Giegling, K. Schönrath, D. Rujescu, Association of VSNL1 with schizophrenia, frontal cortical function, and biological significance for its gene product as a modulator of cAMP levels and neuronal morphology, Transl Psychiatry 1, e22 (2011).

59. J.-M. Lee, K. Blennow, N. Andreasen, O. Laterza, V. Modur, J. Olander, F. Gao, M. Ohlendorf, J. H. Ladenson, The brain injury biomarker VLP-1 is increased in the cerebrospinal fluid of Alzheimer disease patients, Clin. Chem. 54, 1617–1623 (2008).

60. R. Tarawneh, G. D’Angelo, E. Macy, C. Xiong, D. Carter, N. J. Cairns, A. M. Fagan, D. Head, M. A. Mintun, J. H. Ladenson, J.-M. Lee, J. C. Morris, D. M. Holtzman, Visinin-like protein-1: diagnostic and prognostic biomarker in Alzheimer disease, Ann. Neurol. 70, 274–285 (2011).

61. R. Tarawneh, J.-M. Lee, J. H. Ladenson, J. C. Morris, D. M. Holtzman, CSF VILIP-1 predicts rates of cognitive decline in early Alzheimer disease, Neurology 78, 709–719 (2012).

62. C. M. Kirkwood, M. L. MacDonald, T. A. Schempf, A. V. Vatsavayi, M. D. Ikonomovic, J. L. Koppel, Y. Ding, M. Sun, J. K. Kofler, O. L. Lopez, N. A. Yates, R. A. Sweet, Altered Levels of Visinin-Like Protein 1 Correspond to Regional Neuronal Loss in Alzheimer Disease and Frontotemporal Lobar Degeneration, J. Neuropathol. Exp. Neurol. 75, 175–182 (2016).

63. P. M. Keeney, J. Xie, R. A. Capaldi, J. P. Bennett, Parkinson’s disease brain mitochondrial complex I has oxidatively damaged subunits and is functionally impaired and misassembled, J. Neurosci. 26, 5256–5264 (2006).

64. P. Andrés-Benito, E. Gelpi, M. Povedano, G. Santpere, I. Ferrer, Gene Expression Profile in Frontal Cortex in Sporadic Frontotemporal Lobar Degeneration-TDP, J. Neuropathol. Exp. Neurol. 77, 608–627 (2018).

65. D. Martins-de-Souza, W. F. Gattaz, A. Schmitt, C. Rewerts, S. Marangoni, J. C. Novello, G. Maccarrone, C. W. Turck, E. Dias-Neto, Alterations in oligodendrocyte proteins, calcium homeostasis and new potential markers in schizophrenia anterior temporal lobe are revealed by shotgun proteome analysis, J Neural Transm (Vienna) 116, 275–289 (2009).

66. I. S. Piras, C. Bleul, I. Schrauwen, J. Talboom, L. Llaci, M. D. De Both, M. A. Naymik, G. Halliday, C. Bettencourt, J. L. Holton, G. E. Serrano, L. I. Sue, T. G. Beach, N. Stefanova, M. J. Huentelman, Transcriptional profiling of multiple system atrophy cerebellar tissue highlights differences between the parkinsonian and cerebellar sub-types of the disease, Acta Neuropathol Commun 8 (2020), doi: 10.1186/s40478-020-00950-5.

67. R. Tang, H. Liu, Identification of Temporal Characteristic Networks of Peripheral Blood Changes in Alzheimer’s Disease Based on Weighted Gene Co-expression Network Analysis, Front Aging Neurosci 11, 83 (2019).

68. Z. Wang, X. Yan, C. Zhao, Dynamical differential networks and modules inferring disrupted genes associated with the progression of Alzheimer’s disease, Exp Ther Med 14, 2969–2975 (2017).

69. A. Lin, C. J. Giuliano, A. Palladino, K. M. John, C. Abramowicz, M. L. Yuan, E. L. Sausville, D. A. Lukow, L. Liu, A. R. Chait, Z. C. Galluzzo, C. Tucker, J. M. Sheltzer, Off-target toxicity is a common mechanism of action of cancer drugs undergoing clinical trials, Sci Transl Med 11 (2019), doi: 10.1126/scitranslmed.aaw8412.

70. S. Meier, M. Bell, D. N. Lyons, J. Rodriguez-Rivera, A. Ingram, S. N. Fontaine, E. Mechas, J. Chen, B. Wolozin, H. LeVine, H. Zhu, J. F. Abisambra, Pathological Tau Promotes Neuronal Damage by Impairing Ribosomal Function and Decreasing Protein Synthesis, J Neurosci 36, 1001–1007 (2016).

71. Y. Fujiki, A. L. Hubbard, S. Fowler, P. B. Lazarow, Isolation of intracellular membranes by means of sodium carbonate treatment: application to endoplasmic reticulum, J. Cell Biol. 93, 97–102 (1982).

72. M. M. Savitski, T. Mathieson, N. Zinn, G. Sweetman, C. Doce, I. Becher, F. Pachl, B. Kuster, M. Bantscheff, Measuring and managing ratio compression for accurate iTRAQ/TMT quantification, J. Proteome Res. 12, 3586–3598 (2013).

