## Supplemental Materials for "Broad kinase inhibition mitigates early neuronal dysfunction and cognitive deficits in tauopathy"

**This PDF file contains:**

Materials and Methods

Fig. S1. Total duration spent in novel arm is partially rescued in tau transgenic mice treated with GSK2606414.

Fig. S2. Non-PERK UPR proteins do not have altered transcript levels in 5mo tau transgenic mice.

Fig. S3. GSK2606414 treatment for up to 36d does not cause marked weight loss.

Table S1. Top kinase targets of GSK2606414 and relevance to tauopathy.

Table S2. Significantly enriched reactome pathways identified by proteomics.

Table S3. Human to mouse proteomic comparisons.

Table S4. Patient demographics of human proteomic samples.

References *(71, 72)*

**Materials and Methods**

*Tissue Preparation*

Mouse brains were harvested as previously described *(26)*. Briefly, mice were anesthetized using isoflurane and transcardially perfused with saline. The brain was separated into hemispheres where the right hemisphere was drop-fixed in 4% para-formaldehyde for immunohistochemistry and the left hemisphere was microdissected, snap frozen in liquid nitrogen, and stored at -80°C until homogenization. For immunoblot analyses, brains were weighed and diluted in homogenization buffer (1:100 Protease Inhibitor, Calbiochem; 1:100 Phosphatase Arrest II, G Biosciences; 1:100 Phosphatase Arrest III G Biosciences; 100µM PMSF; in 1x RIPA, Thermo) was added to a final concentration of 10% (w/v). Samples were homogenized using a using a small electronic pestle (VWR) and centrifuged at 4°C at 13,000g for 15 minutes. Protein concentration was established using the bicinchoninic acid (BCA) assay (ThermoScientific).

*Protein analysis of brain homogenates*

Western blots were performed as previously described *(26)*. Briefly, proteins were separated via SDS-PAGE, transferred to PVDF membranes, blocked in 5% (w/v) milk-TBS for 1 hour, and then incubated in primary antibody overnight at 4°C. Primary antibodies diluted in 5% (w/v) milk-TBS: PHF1 1:1000 (Peter Davies), H150 total tau 1:1000 (SantaCruz), pY216-GSK3b 1:1000 (Abcam), total GSK3b 1:1000 (CST), 3-NT (1:500, Santa Cruz),and GAPDH and actin for loading controls (1:1000, CST). Different loading controls were used throughout experiments to ensure optimal normalization given non-specific antibody binding and species reactivity. HRP-conjugated secondary antibodies to mouse or rabbit IgG (Southern Biotech and ThermoScientific) were used at a 1:1000 dilution where appropriate. Bands were visualized using ECL (ThermoScientific). Image quantification was performed using ImageJ, normalized to loading controls.

*Open Field Test*

Open field-testing was performed in the University of Kentucky Rodent Behavior Core. All testing was performed at the same time of day for each cohort. Animals were acclimated to the testing room 30 minutes prior to testing. Open field-testing was done with the automated Photobeam Activity System (PAS) with Flexfield Animal Activity System (San Diego Instruments, Inc. San Diego) coupled to a computer used to eliminate human interaction and bias. Each animal was placed into the middle of the open field, facing away from the experimenter and exploratory behavior recorded for 30min. The chamber was cleaned with 10% ethanol between tests to eliminate olfactory cues from previous animals. The time mice spent in the peripheral areas were recorded in six, 5-min intervals and is reported by normalizing to time spent in the first 5-min window. Recordings were taken using EthoVision XT and reviewed by two blinded investigators. Several cohorts were tested independently over periods of months, but each animal was tested on the 21^st^ day of treatment.

*Y-maze*

Forced alternation test was performed according to a previously published protocol (*30*). A symmetrical Y-maze (Stoelting Co, Wood Dale, IL, USA) with grey Perspex® walls and steel bottom (arms: 35 cm long, 5 cm wide, 10 cm high) was used for the test. The light in the testing area was kept at 30 ± 5 lux and the testing room was maintained at room temperature (23 ± 2°C). Mice were handled for 5 days (5min/day/mouse) before testing. During the 5 min sample trial (T1), mice explored two arms of the Y-maze with the third arm blocked. The mice were placed into the end of the start arm facing the wall, away from the center. After completion of the sample trial, each mouse was returned to its home cage for 30 min. During the test trial (T2), the block in arm 3 was removed and the mouse was allowed to explore the entire maze for 5 min. In between animals and trials, the maze was cleaned with Quatricide® solution to prevent odor cues.

*Manganese-enhanced magnetic resonance imaging (MEMRI)*

All MRI procedures were performed in accordance with the University of Kentucky MRISC IACUC Protocol. Brain volume was calculated from T2 images based on manual tracing of the brain. Manganese chloride (30mM) prepared in saline was delivered to mice via intraperitoneal injection (66 mg/kg). All MR imaging was performed on a horizontal bore 7T Bruker ClinScan magnet (Bruker, Ettlingen, Germany) using a cylindrical volume coil for excitation and a cryocoil for detection. In preparation for imaging, animals are anesthetized using isoflurane (3-5%) in oxygen at a rate of 0.5-1.0 L/min and maintained using 1-3% isoflurane in oxygen. Body temperature was maintained using circulating water, and vital signs (core temperature and respirations) were monitored using a physiological monitoring system SA Instruments Inc. (SAI, Stony Brook, New York). When applicable, MEMRI was performed after behavioral testing to reduce stress-related confounds in behavioral test results.

Look-locker imaging was performed following non-selective spin inversion in one slice of the hippocampus. Fifty images were acquired following inversion with image spacing of 100ms (total sequence repetition time of 5s) to fully sample the T1-relaxation curve. Additional image parameters included TR/TE = 5500/1.9, Matrix = 128 x 128, number of averages = 3, field of view = 17mm x 17mm x 0.7mm. T2-weighted images were acquired covering the entire brain (excluding the cerebellum and olfactory bulb) using a turbo-spin echo sequence with TR/TE = 3360/42, Slices = 21, Matrix = 448 x 336, number of averages = 2, field of view = 25mm x 25mm x 0.5mm. The imaging procedures for scanning one mouse were completed in 45 min. Imaging was performed prior to the injection of MnCl_2_ (baseline) and repeated at 6h post-injection. Image mapping and analysis was performed in MATLAB (Mathworks, Nattic, MA, USA). Images from the Look-Locker series were used to reconstruct voxel-by-voxel signal relaxation curves which were fit to the equation S(t) = So(1-2*exp(-R1*T1)), where S(t) represents the signal at a given inversion time (TI), So represents the steady state signal at maximal T1, and R1 represents the longitudinal relaxation rate. Regions of interest (DG, CA1, CA1, CA3, and superior medial cortex) were identified using the Allen Brain Mouse Atlas. Within each the change in R1 relaxation rates (ΔR1) before and after MnCl_2_ exposure was calculated as:

$$\Delta R_{1}={R1}_{6h}-{R1}_{baseline}$$

*Quantitative real-time RT-PCR gene card arrays*

Total RNA was extracted from snap frozen hippocampal tissues using TRI reagent (Ambion) following the manufacturer’s protocol. RNA was quantified using a BioTech spectrophotometer and cDNA was generated using SuperScript IV reverse transcriptase (Invitrogen). Real-time RT-PCR was performed using a TaqMan custom Array (Life Technologies) according to manufacturer’s instructions. Gene expression was evaluated by normalizing to GAPDH and 18S expression as internal controls. The real-time values for each sample were averaged and evaluated using the comparative CT (cycle time) method.

*Immunofluorescence*

Immunofluorescence (IF) was performed as previously described *(26, 70)*. Primary antibodies: pPERK (1:250, SantaCruz) and NRF2 (1:250; Abcam).

*Protein Extraction and Quantification for Proteomics*

Microdissected hippocampal mouse tissue were processed for protein extraction as described previously *(71)* with the following modifications. Briefly, samples were ground in liquid nitrogen into fine powder and incubated in extraction buffer (0.1 M Tris-HCl pH 8.8, 10 mM EDTA, 0.2 M DTT, 0.9 M sucrose), ground in a fume hood, and the resulting extract was agitated for 2 hrs at room temperature. After washing twice with 0.1 M ammonium acetate in methanol and twice with 80% acetone, the dried pellet was dissolved with 50mM ammonium bicarbonate buffer. The mixtures were cooled on ice before centrifugation (4°C) for 20 min at 12,000 g. The soluble proteins were performed using an EZQ Protein Quantitation Kit (Invitrogen, Carlsbad, CA, USA) with the SoftMax Pro Software v5.3 (Molecular Devices, Downingtown, PA, USA).

*Protein Extraction, Digestion, TMT Labeling, and LC-MS/MS*

Proteins were dissolved in protein buffer (supplemental materials) (8M Urea, 50 mM Tris-HCl (pH 8.0), 50 mM triethylammonium bicarbonate, 0.1% SDS (w/v), 1% Triton-100 (w/v), 1 mM PMSF, 10 μg/ml leupeptin, 1% phosphatase inhibitor cocktail 2 and 3 (v/v)). For each sample, a total of 50 μg of protein were reduced, alkylated, trypsin-digested, and labeled according to the manufacturer’s instructions (Thermo Scientific). For each mass-spectrometer batch, late-stage 8mo transgenic tissue were labeled with TMT tags 126 and 127N, non-transgenic mice treated with 414 were labeled with TMT tags 128C and 129N, non-transgenic mice treated with vehicle were labeled with TMT tags 128C and 129N, transgenic mice treated with 414 were labeled with 129C and 130N, transgenic mice treated with vehicle were labeled with 130C and 131N, and a mixed-sample internal batch was labeled with 131C. Labeled peptides were desalted with C18-solid phase extraction and dissolved in strong cation exchange (SCX) solvent A (25% (v/v) acetonitrile, 10 mM ammonium formate, and 0.1% (v/v) formic acid, pH 2.8). The peptides were fractionated using an Agilent HPLC 1260 with a polysulfoethyl A column (2.1 × 100 mm, 5 µm, 300 Å; PolyLC, Columbia, MD, USA). Peptides were eluted with a linear gradient of 0–20% solvent B (25% (v/v) acetonitrile and 500 mM ammonium formate, pH 6.8) over 50 min followed by ramping up to 100% solvent B in 5 min. The absorbance at 280 nm was monitored and a total of 19 fractions were collected. The fractions were lyophilized, desalted, and resuspended in LC solvent A (0.1% formic acid in 97% water (v/v), 3% acetonitrile (v/v)). A hybrid quadrupole Orbitrap (Q Exactive) MS system (Thermo Fisher Scientific, Bremen, Germany) was used with high energy collision dissociation (HCD) in each MS and MS/MS cycle. The MS system was interfaced with an automated Easy-nLC 1000 system (Thermo Fisher Scientific, Bremen, Germany). Each sample fraction was loaded onto an Acclaim Pepmap 100 pre-column (20 mm × 75 μm; 3 μm-C18) and separated on a PepMap RSLC analytical column (250 mm × 75 μm; 2 μm-C18) at a flow rate at 350 nl/min during a linear gradient from solvent A (0.1% formic acid (v/v)) to 30% solvent B (0.1% formic acid (v/v) and 99.9% acetonitrile (v/v)) for 95 min, to 98% solvent B for 15 min, and hold 98% solvent B for additional 30 min. Full MS scans were acquired in the Orbitrap mass analyzer over m/z 400–2000 range with resolution 70,000 at 200 m/z. The top ten most intense peaks with charge state ≥ 3 were fragmented in the HCD collision cell normalized collision energy of 28%, (the isolation window was 2 m/z). The maximum ion injection times for the survey scan and the MS/MS scans were 250 ms, respectively and the ion target values were set to 3e6 and 1e6, respectively. Selected sequenced ions were dynamically excluded for 60 sec.

Raw data files obtained from the Orbitrap Fusion were processed using Proteome Discoverer™ (version 2.3). MS/MS spectra were searched against the UniProt mouse proteome database downloaded Dec 4, 2019 with 63,369 total sequences including the sequence for 0N4R P301L tau mutant cDNA. The SEQUEST HT search engine was used and parameters were the following: fully-tryptic specificity; maximum of two missed cleavages; minimum peptide length of 6; fixed modifications for TMT tags on lysine residues and peptide N-termini (+ 229.162932 Da) and carbamidomethylation of cysteine residues (+ 57.02146 Da); variable modifications for oxidation of methionine residues (+ 15.99492 Da); phosphorylation of serine, threonine and tyrosine (+ 79.9663 Da); precursor mass tolerance of 20 ppm; fragment mass tolerance of 0.05 Daltons. Percolator was used to filter peptide spectral matches (PSM) and peptides to a false discovery rate (FDR) of less than 1% using target-decoy strategy. Following spectral assignment, peptides were assembled into proteins and were further filtered based on the combined probabilities of their constituent peptides to a final FDR of 1%. In cases of redundancy, shared peptides were assigned to the protein sequence in adherence with the principles of parsimony. Reporter ions were quantified from MS2 scans using an integration tolerance of 20 ppm with the most confident centroid setting.

Each MS batch protein abundance value was normalized to the internal standard (131C) abundance, enabling batch-corrected analysis across groups. The criteria for protein inclusion in statistical analysis were that each protein must i) be present in all samples, ii) have a global FDR < 0.05, iii) contain at least one unique peptide, and iv) be an annotated master protein. These filtering criteria reduced the total amount of unique protein hits to 24,149 peptides and 1100 proteins. Normalized abundance ratios of selected proteins were loaded into R (3.6.2) for processing. Significant differential expression of proteins between samples was determined as p < 0.05 using unpaired Student’s t-test with FDR multiple-comparisons correction. No fold change (FC) cutoff was used as TMT-labeled tagging and subsequent analysis is subject to ratio compression *(72)*. Reactome pathway *(32)* changes were assessed using gProfiler *(31)* were used against total mouse protein coding background. Significant Reactome pathways (FDR adj. p value < 0.10) were loaded into Cytoscape for clustering analysis *(33)*. For comparison to published human proteomic data, Student’s t-test with FDR or Tukey correction was used to compare between diseased groups and control samples.

*IMAC-enriched phospho-proteomics*

Phosphopeptide Enrichment is performed using titanium dioxide and zirconium dioxide Nutip (Glygen Inc., Columbia, MD, USA). First, the Nutip is equilibrated by washing the resin 10X with 5 µl of binding solution (80% acetonitrile (v/v), 5% TFA (v/v), pH 3), and the tryptic digests were mixed 1:1 with binding solution. The mixture was loaded onto the Nutip by taking 2.5 µl aliquots and aspirating and expelling each aliquot 50 times over 30 minutes. Washes consisting of 80% acetonitrile (v/v), 1% TFA (v/v) (pH 3) followed 10 times. The bound phospho-peptides were eluted with 10 µl of elution solution (2% NH4OH (v/v), pH 11). To prevent loss of the phosphate moieties under highly basic conditions, the eluent was acidified with 2 µl of 10 % formic acid (v/v) and lyophilized. The sample was resuspended in LC solvent A and injected into the Q Exactive Plus (see above).

**Supplementary Figures and Tables:**

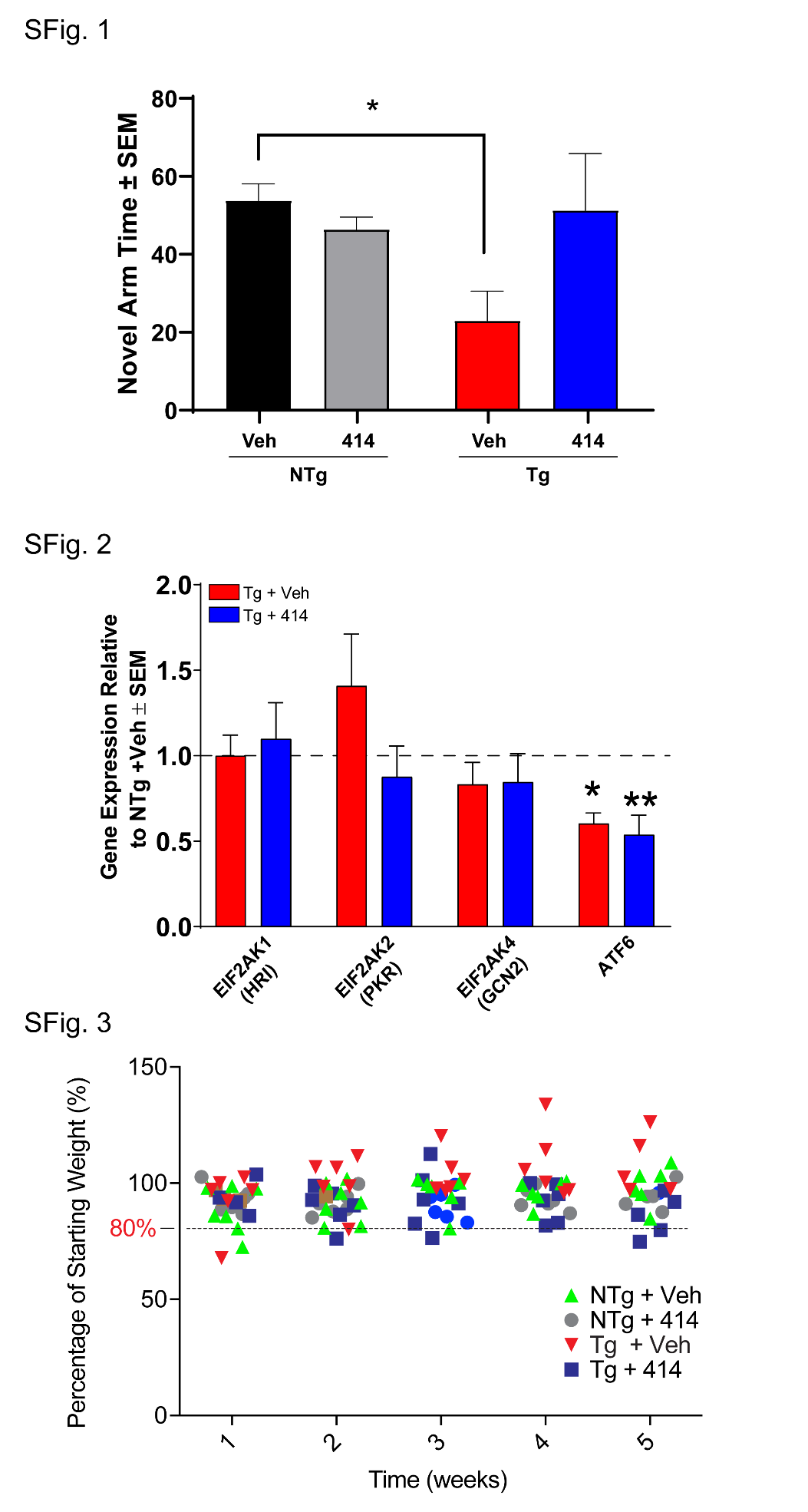

**Fig. S1. Total duration spent in novel arm is partially rescued in tau transgenic mice treated with GSK2606414.** Total time was measured during mouse exploration in the Y-maze behavioral task. Tg + Veh mice exhibited reduced time spent in the novel arm (*p = 0.0433). Tg mice treated with 414 were only partially rescued compared to Tg + Veh mice (p = 0.0882). Data were compared using two-way ANOVA with Tukey *post hoc* test.

**Fig. S2. Non-PERK UPR proteins do not have altered transcript levels in 5mo tau transgenic mice.** Other eIF2a kinases do not have altered transcript levels in 5mo Tg mice. *ATF6* transcript levels decrease in Tg mice compared to NTg controls, though 414 has no effect. Transcript levels normalized to GAPDH and 18S (n = 4-6). Two-way ANOVA with Tukey *post hoc* test. Data are expressed as the mean relative to NTg + Veh ± SEM, *p < 0.05, **p < 0.01.

**Fig. S3. GSK2606414 treatment for up to 36d does not cause marked weight loss.** Treatment with 100 mg/kg GSK2606414 treatment for 30 – 36d did not induce significant weight loss.

**Table S1. Top kinase targets of GSK2606414 and relevance to tauopathy.** The top targets of GSK2606414 with experimentally determined IC50 are listed based on reported data *in vitro (10)* or based on cell culture experiments *(17, 18)*.

**Table S2. Significantly enriched reactome pathways identified by proteomics.** Reactome pathways significantly altered (FDR adjusted p < 0.10) by either direction in Tg mice compared to NTg controls along with pathways determined rescued by 414.

**Table S3. Human to mouse proteomic comparisons.** Protein abundances shared between mouse and human datasets and related statistical analyses.

**Table S4. Patient demographics of human proteomic samples.** Patient demographics and disease-related pathology scores were averaged and compiled from original reports *(37, 39)*.
